# Calibration-free molecular counting from a single DNA-PAINT intensity trace using cumulants

**DOI:** 10.1101/2025.11.17.688450

**Authors:** Teun A.P.M. Huijben, Rodolphe Marie, Jonas N. Pedersen

## Abstract

Single-molecule localization microscopy achieves nanometer-scale resolution but fails to count molecular targets when multiple targets are in close proximity. DNA-PAINT uses reversible binding of fluorescently labeled probes to image molecular targets, but existing time-series based counting methods require prior knowledge of binding kinetics, calibration, or extensive data post-processing, which limits applicability across heterogeneous biological samples. Here, we present mCOAST, a calibration-free method that extracts molecular counts and kinetic parameters directly from a single DNA-PAINT intensity trace using cumulants. Unlike existing approaches, mCOAST requires no adjustable parameters, data normalization, or denoising. We demonstrate accurate counting for diffraction-limited clusters with up to 48 targets at high imager concentrations, with precision that improves, rather than decreases, as concentration increases. Critically, mCOAST counts targets in individual clusters despite kinetic heterogeneity across samples or between experiments. This paves the way towards quantitative imaging and counting in uncalibrated biological systems, such as living cells.

## Introduction

Super-resolution microscopy has drastically changed the fields of biological and material sciences by visualizing structures at the nanometer scale [1]. Single-molecule localization microscopy methods, a subset of super-resolution approaches, circumvent the diffraction limit by using blinking fluorescent dyes, where repeated imaging of sparse subsets of fluorophore-labeled targets ensures that each fluorophore creates a single isolated diffraction-limited spot on the camera [2, 3, 4]. The position of each fluorophore is determined with nanometer precision by fitting the fluorescent spots in each frame [5]. This allows, in principle, for counting nanometer-sized target molecules by simple visual inspection of the super-resolved image, but the length of the linker between the fluorophore and the target of interest, sample drift, and molecular motion introduce localization errors, resulting in a single cluster of localizations. Recent sub-nanometer resolution techniques like MINFLUX [6] and RESI [7] would, in principle, allow visual molecular counting, but they require perfectly fixed samples and known structures.

An alternative approach to the counting problem is to utilize that each target in a cluster contributes a number of blinking events. Multiple approaches based on PALM and STORM demonstrate molecular counting by calibrating the number of localizations per fluorophore [8, 9, 10, 11], the distribution of dark-times [12], an assumed model for the photophysics of the fluorophore [13, 14, 15, 16, 17, 18], or spatial clustering of the localizations [19, 20]. All these approaches are limited by the complex photophysics of the blinking dyes. In a DNA-PAINT experiment, the blinking is instead the result of binding and unbinding of the fluorophore to the molecular target, facilitated by DNA-hybridization [21]. Single-stranded DNA probes (‘docking strands’) are chemically conjugated to the molecule of interest, to which dye-labeled complementary imager strands (‘imagers’) transiently bind (Fig. 1**b**). The docking strand, referred to as a ‘target’, is in an ‘on’-state when an imager is bound and otherwise in an ‘off’-state. DNA-PAINT has proven to be the method of choice for molecular counting since the hybridization kinetics are governed by easy-to-model chemical reactions, and the kinetic parameters can be experimentally altered by changing either the sequence of the DNA strands or the temperature and ionic strength of the buffer. Furthermore, the intrinsic dye-exchange mechanism provides robustness against photobleaching of fluorophores.

**Figure 1:**
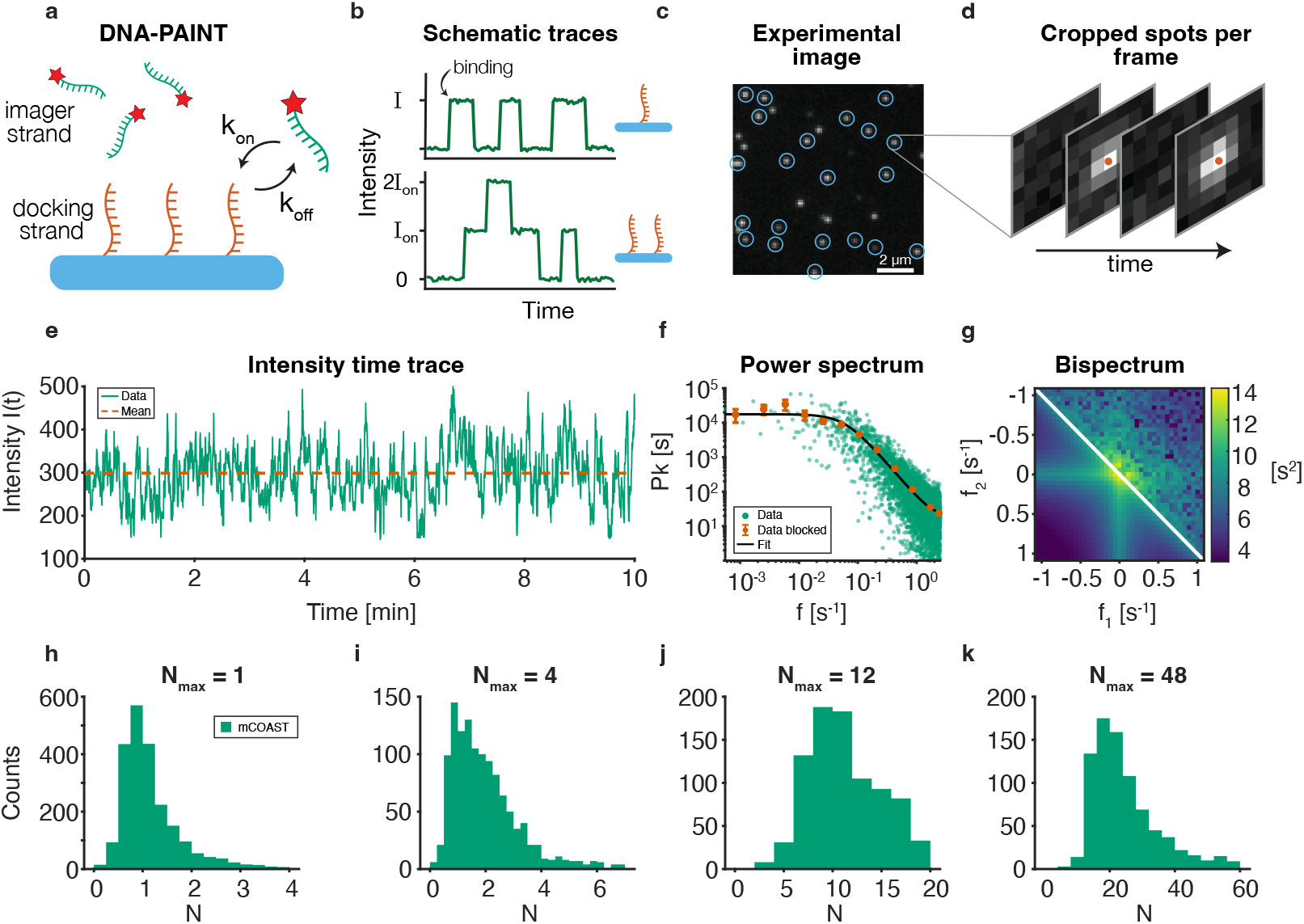
Calibration-free molecular counting of individual molecular clusters with mCOAST. **a** The object of interest (blue) is functionalized with single-stranded DNA docking strands (red), to which dye-labeled (red star) complementary single-stranded DNA imager strands (green) transiently bind, governed by binding and unbinding rates 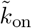 and *k*_off_ . **b** Every binding event results in an increase in intensity, reflected in the intensity time trace. For a single docking strand (*N* = 1, top trace), the binding events are separated in time because they happen on the same docking strand, while for multiple closely-spaced docking strands (*N* = 2, bottom trace), the binding events can occur simultaneously. **c** For experimental data, individual localizations are grouped into spots from individual clusters. **d** For each cluster, we record the position (red dots), the spot intensity, and the background intensity at every time point. The intensity time trace used for analysis is calculated by subtracting the background intensity from the spot intensity. **e** Sample of an experimental trace, including the mean value (dashed red line). **f** Experimental power spectrum with fit to Eq. (7). **g** Experimental bispectrum (upper half) with fit to Eq. (8) (lower half). **h-k** Molecular counting with mCOAST for individual experimental traces for *N*_max_ = 1, 2, 4, and 48 and imager concentration *c*_i_ = 20 nM. The number of traces is stated in Supplementary Note 1, Table S1.

Quantitative PAINT (qPAINT [22]) provides molecular counting for individual clusters of molecules based on the distribution of dark-times, i.e., time intervals with no fluorophores bound to any target. Despite its simplicity and robustness, the major drawback of qPAINT is that the molar binding rate must be known from a calibration experiment as it depends strongly on experimental conditions, e.g., temperature [23, 21, 24, 25, 26], buffer conditions [24, 26], and steric hindrance from the direct surroundings of the docking strands. Importantly, the imager concentration must also be tuned in each experiment to ensure the presence of dark-times. With localization-based fluorescence correlation spectroscopy (lbFCS [26]), absolute molecular counting is performed based on time series analysis. lbFCS does not require knowledge of the binding rate, but it still requires multiple calibration experiments and assumes that the binding kinetics are identical between experiments and among all clusters. Calibration-free alternatives that overcome the need for a priori knowledge of the binding kinetics have also been developed. An updated approach of lbFCS, lbFCS+ [27], performs molecular counting and extraction of hybridization kinetics for individual clusters of molecules, but it relies on a normalization of the intensity traces, which is not possible for high background intensities and experimentally realistic noise levels. lbFCS+ only works for a small number of molecules (*N* < 10), and requires tuning of the imager concentration.

Here, we present molecular counting from a single time trace (mCOAST) from a DNA-PAINT experiment without any prior knowledge or sensitive data processing. First, we model a DNA-PAINT intensity trace mathematically as a sum of identical and independent two-state jump processes and show how to express the number of targets and the kinetic parameters in terms of only the first three cumulants of the trace and a characteristic time scale [Eqs. (1-3) and Fig. 1]. The cumulants are extracted from fits to the power- and the bispectra [Eqs. (4,5) and Fig. 1**e,f**]. We demonstrate the method on experimental data for DNA origami structures with varying numbers of docking strands and at different imager concentrations, where we obtain the number of targets in the clusters (Fig. 1**h-k**) and the kinetic parameters (Fig. 2). Using ensemble-averaged kinetic parameters for clusters within the same field of view increases precision on the estimated number of targets in individual clusters, and extends the range of the method to lower imager concentrations (Fig. 3). Uncertainties on the estimated parameters and the influence of experimental noise is investigated with numerical simulations for experimentally relevant parameters (Fig. 4). A detailed derivation of the main results is in Supplementary Note 2.

**Figure 2:**
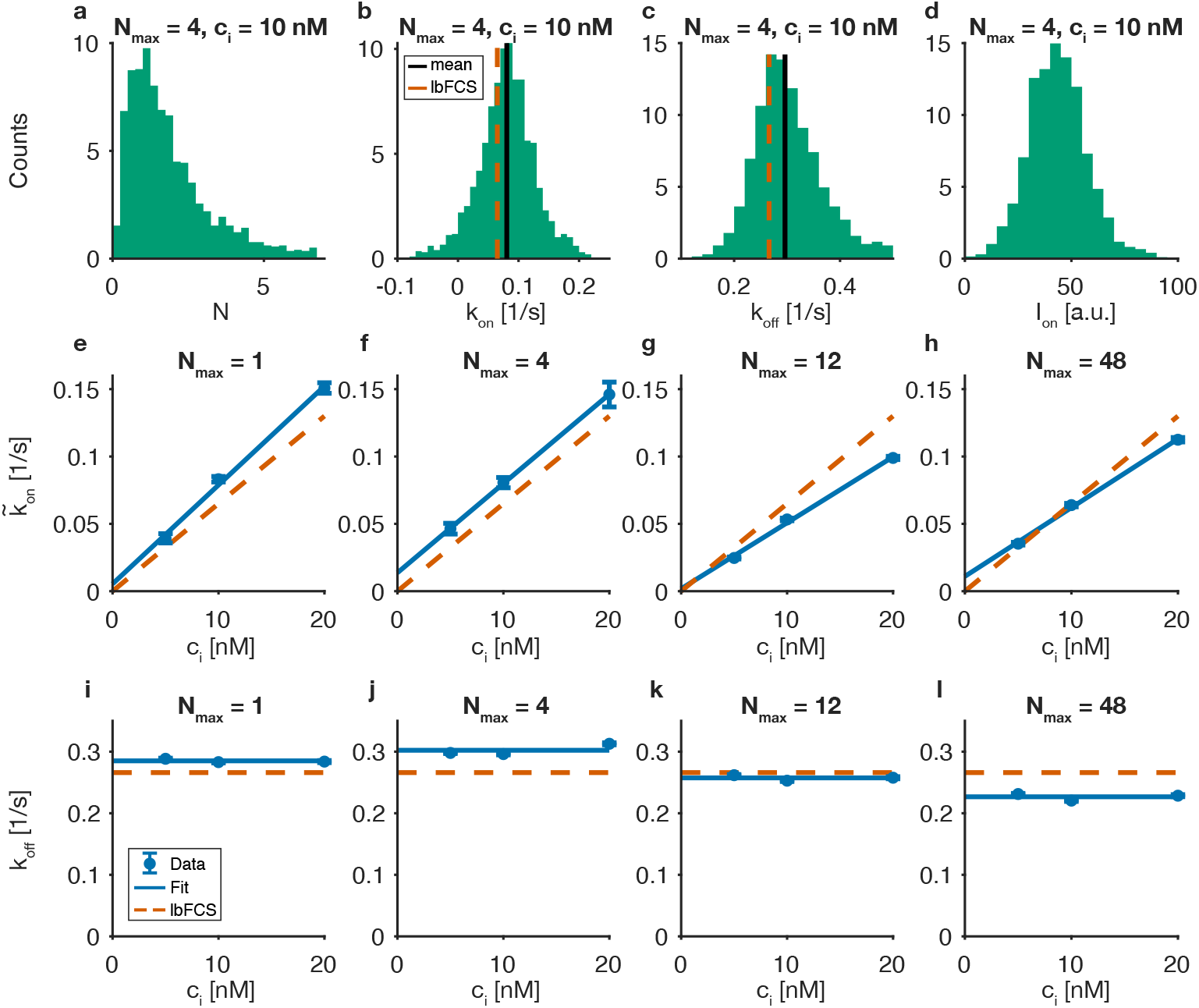
Extraction of DNA binding kinetics with mCOAST. **a-d** Distribution of the four estimated parameters (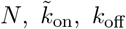, and *I*_on_) by mCOAST for *N*_max_ = 4 origami structures imaged with an imager concentration *c*_i_ = 10 nM. In Panels **b**,**c**, solid, vertical blue lines show the mean, and the dashed, vertical red lines are the result from lbFCS [26]. **e-h** Mean of estimated 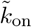 values for four different origami designs as a function of imager concentration. Error bars are standard errors on the means. The solid, blue line is a fit to the data. The slopes correspond to *k*_on_ and the dashed, red line is the result from lbFCS with *k*_on_ = 6.5 · 10^6^ M^−1^s^−1^[26]. **i-l** Means of estimated *k*_off_ values for the four different origami designs as a function of imager concentration. Error bars are standard errors on the means. The solid, blue line is the weighted average, and the dashed, red line is the result from lbFCS with *k*_off_ = 0.266 s^−1^[26]. The number of traces per condition is stated in Supplementary Note 1, Table S1.

**Figure 3:**
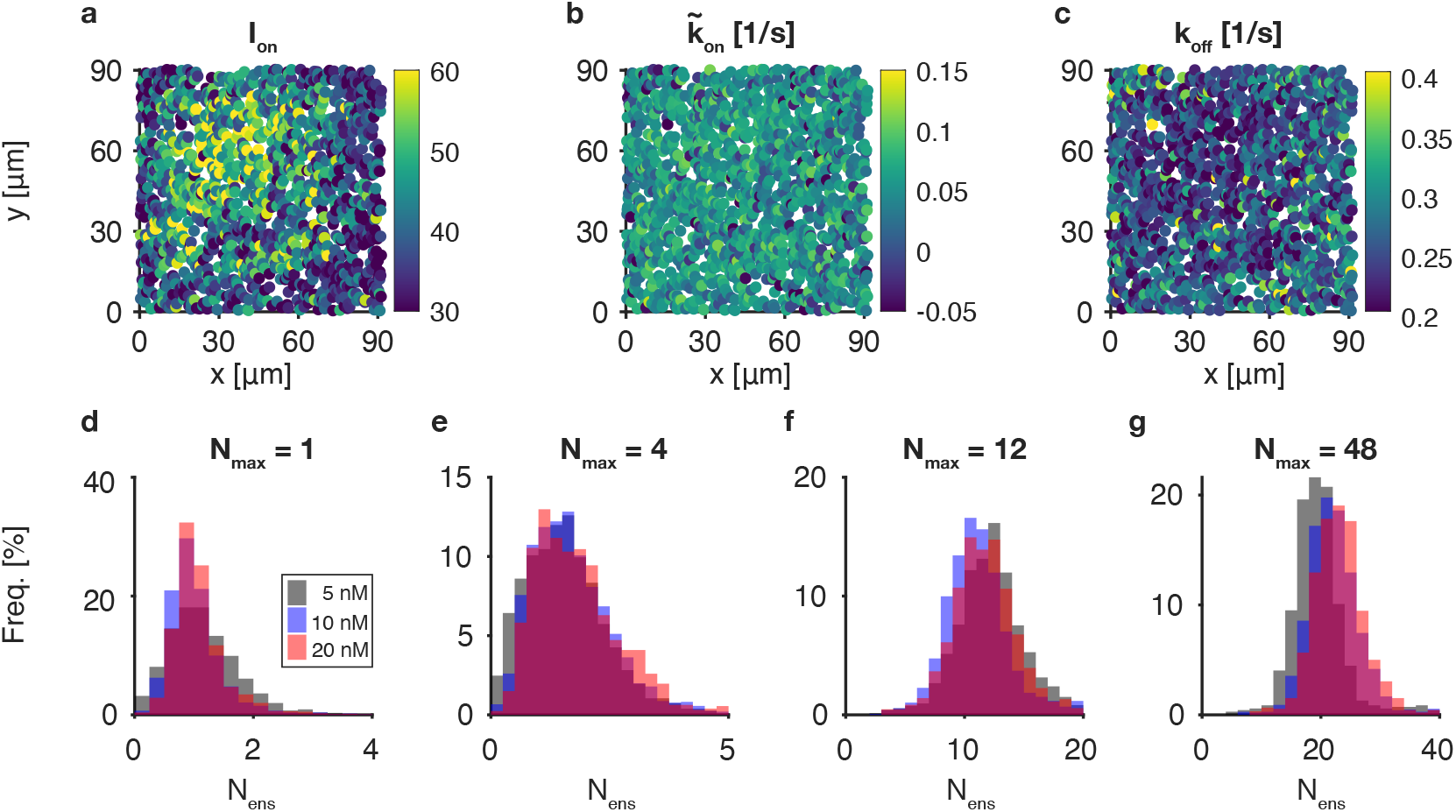
Ensemble-averaging kinetic parameters for spots in the same field of view. Panels **a-c** show the estimated values for *I*_on_, 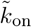, and *k*_off_ across the entire field of view for *N*_max_ = 12 and *c*_i_ = 10 nM. The rates 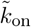 and *k*_off_ exhibit no significant spatial variation, while the highest values of *I*_on_ are concentrated in the upper left corner, highlighting the non-uniform excitation illumination pattern. Panels **d-g** are the estimates for the number of targets from Eq. (6) with the average kinetic rates as input for all four origami structures and all three imager concentrations. The number of traces is stated in Supplementary Note 1, Table S1.

**Figure 4:**
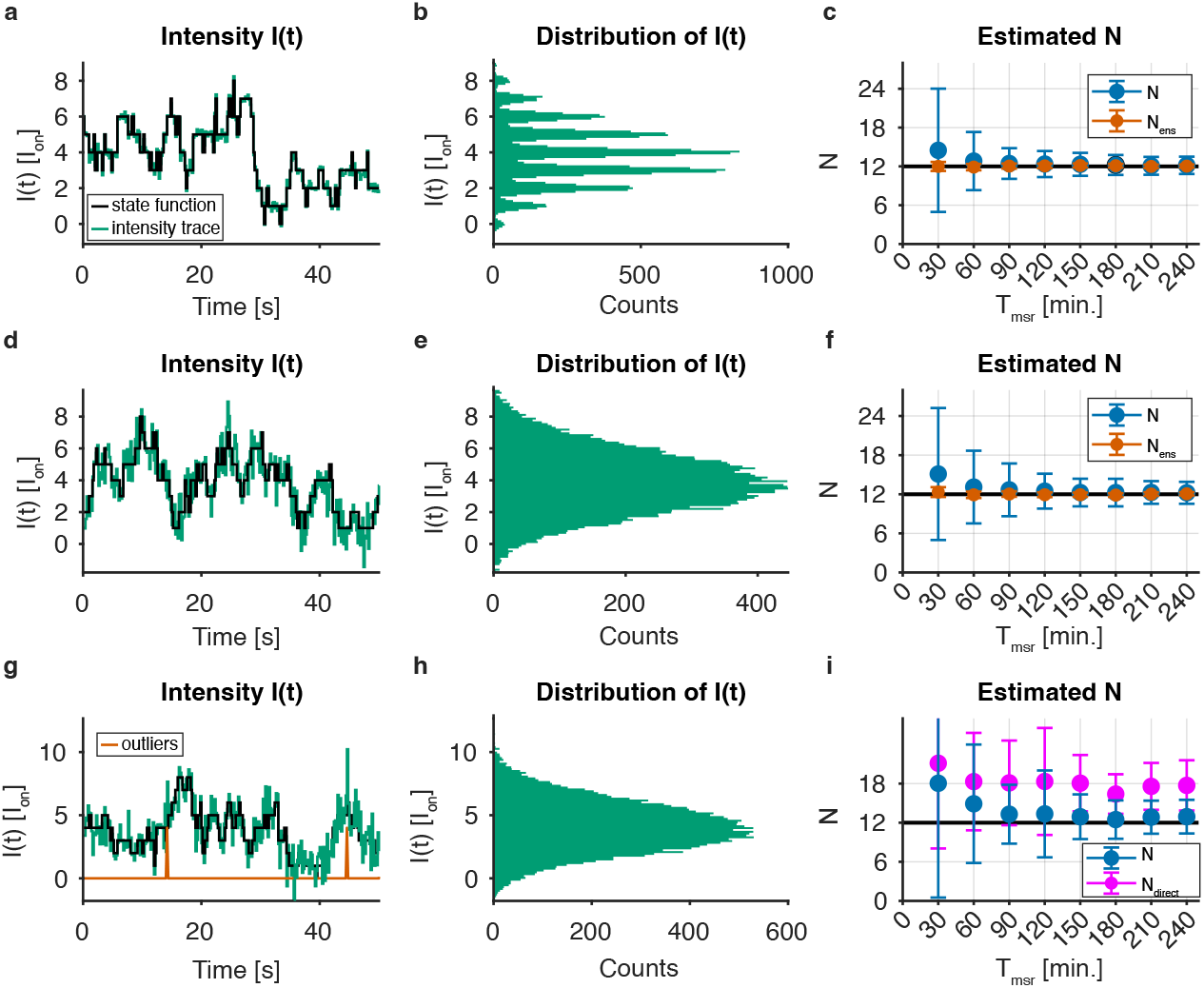
Results of mCOAST on simulated trajectories. DNA origami (black) and intensity traces *I*(*t*) (green) for *σ*_bg_ = 0.2*I*_on_ and *N* ^(true)^ = 12, **b** corresponding histogram of intensity values for a measurement time *T*_msr_ = 60 min, and **c** the estimated number of targets *N* [Eq. (1)] and *N*_ens_ [Eq. (6)] versus measurement time. **d-f** Same as panels **a-c**, except *σ*_bg_ = *I*_on_. **g-i** Same as panels **d-f**, except that Poisson distributed outliers with a rate *µ*_outlier_ = 0.03*/*s and amplitude *I*_outlier_ = 4*I*_on_ have been added to the signal (red spikes in panel **g**). Estimates *N*_direct_ are from a direct calculation of the cumulants from the intensity trace. For all panels: *k*_on_ = 6.5 · 10^6^ M^−1^s^−1^, *k*_off_ = 0.266 s^−1^, *c*_i_ = 20 nM, and Δ*t* = 0.2 s. Simulations are repeated 1000 times for all parameter sets. Solid dots are mean values, and error bars are standard deviations.

## Results

### Mathematical model of a DNA-PAINT intensity trace

Similar to all alternative approaches for molecular counting based on time-series analysis, we assume that the four parameters describing a DNA-PAINT intensity trace from a molecular cluster are the effective association rate 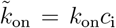 (*c*_i_ is the imager concentration), the dissociation rate *k*_off_ for a single target, the intensity of a single binding event *I*_on_ (Fig. 1**a**), and the number of targets *N* . It is also assumed that all targets within the molecular cluster give the same observed intensity *I*_on_ and have the same binding kinetics (identical), that binding is not affected by neighboring targets (independent), and that the number of targets *N* and the binding kinetics are constant over the course of the experiment (time-translation invariance). So, a molecular cluster with *N* targets can have any integer number between 0 and *N* imagers attached, and each target produces a signal *I*_on_ when in the ‘on’-state, and zero signal otherwise. The binding and unbinding of imagers to the molecular cluster produce a time-dependent intensity trace *I*(*t*). Due to the constant rates, the waiting times for a single site being in an off (on) state are exponentially distributed with a characteristic time 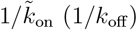 [22]. The average time of a binding-unbinding event is 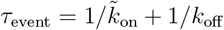, and the expected number of events for each target during an experiment with measurement time *T*_msr_ is *n*_event_ = *T*_msr_*/τ*_event_.

With the assumptions made, a DNA-PAINT intensity trace can be described mathematically as a sum of independent two-state jump processes, one for each target in the molecular cluster (Fig. 1**b**). The two-state jump process, also known as a telegraph process or a dichotomous random process [28], is a Markovian continuous-time stochastic process that also appears, e.g., in single ion-channels [29], current noise in semiconductors [30], and barrier-tunneling in solid-state physics [31].

### Model parameters from the cumulants of a DNA-PAINT intensity trace

Cumulants (also named semi-invariants) are mathematical objects with the property that the cumulant of the sum of independent variables is the sum of their cumulants [32]. Due to the assumption of identical and independent targets and the additive property of cumulants, the first three cumulants of an intensity trace from a cluster with *N* targets are 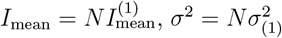 and 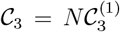, where the first three cumulants of an intensity trace with a single target are the mean 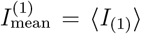, the variance 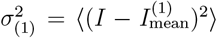, and the third-order central moment 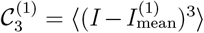. For the two-state jump process, the cumulants can be expressed in terms of the model parameters as 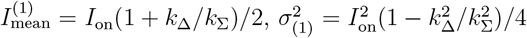, and 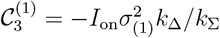, with 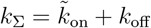 and 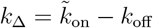 (see Supplementary Note 2). Solving for the number of targets gives

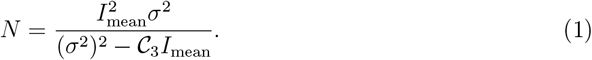

That is, the number of targets in a cluster is expressed solely by the first three cumulants of the intensity trace, which is the main result of the paper. Solving for the kinetic parameters gives that they, next to the first three cumulants, also depend on the sum of the rate *k*_Σ_,

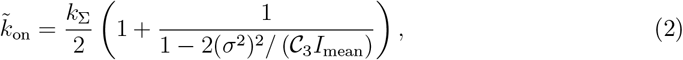

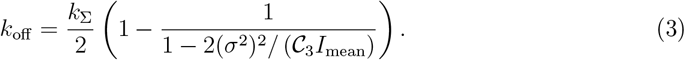

Likewise, concluding our full set of equations, the intensity of a single binding event is *I*_on_ = 2*σ*^2^*/I*_mean_ − C_3_*/σ*^2^.

### Cumulants from experimental intensity traces

The procedure for extracting the experimental intensity time traces for a single cluster and the background from images is outlined in Fig. 1**c** (see also Supplementary Note 1, Fig. S1). Images are recorded with a constant frame rate with a sample time Δ*t*, and from the background-subtracted intensity in frame *j, I*_*j*_ = *I*(*j*Δ*t*) (*j* = 1, 2, … *n*), we calculate the mean intensity 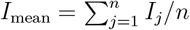. The variance *σ*^2^ and the third-order cumulant 𝒞_3_ can, in principle, be estimated directly from the recorded signal as 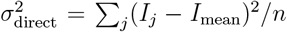 and 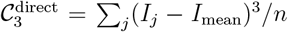. The intensity from a cluster contains, however, both the intensity from the imager strands attached to the docking strands but also contributions from, e.g., the fluorescence from freely diffusing imager strands and the isolated intensity peaks, so-called outliers. The outliers arise, e.g., from diffusing objects that pass close by the cluster and create high-intensity values that only last a single frame. So, extracting third- or higher-order cumulants directly from data should be done with caution [33].

Instead, we reduce the effects of both background intensity and outliers by estimating *σ*^2^ and 𝒞 _3_ from fits to the power spectrum *P* (*ω*) and the bispectrum *B*(*ω*_1_, *ω*_2_) of the intensity trace. The power spectrum and the bispectrum are the Fourier transforms of the second- and third order cumulant sequences, i.e., *C*_2_(*t, t*^′^) = ⟨ [*I*(*t*) −⟨*I*⟩ ][*I*(*t*^′^) −⟨*I*⟩ ] ⟩ and *C*_3_(*j, j*^′^, *j*^′′^) = ⟨ (*I*_*j*_ − *I*_mean_)(*I*_*j*_*′* − *I*_mean_)(*I*_*j*_*′′* −*I*_mean_) ⟩ [34]. Like cumulants, the cumulant sequences have the property that the cumulant sequence of a sum of independent processes is the sum of the cumulant sequences for the individual processes [35].

For a time-translationally invariant two-state jump process, the power spectrum is given by (Supplementary Note 2) [30]

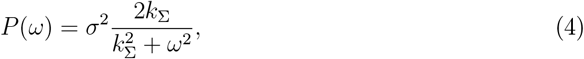

and the bispectrum is (Supplementary Note 2)

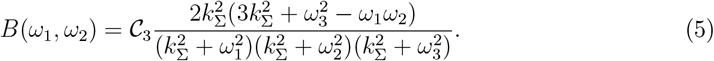

Here *ω* is the angular frequency and *ω*_3_ = *ω*_1_ + *ω*_2_. We fit the experimental power- and the bispectrum to Eqs. (7,8), which are modified versions of Eqs. (4,5) that take into account the discrete nature of the sampled data, and are corrected for aliasing artifacts (see Methods). An example of an intensity trace and fits to the power- and bispectra are shown in Fig. 1**e-g**. The fit to the power spectrum gives both the variance (*σ*^2^) and the sum of the rates (*k*_Σ_). A Gaussian white-noise background intensity gives a constant background in the power spectrum, which is included in the fit, but as cumulants of order greater than two vanish for a Gaussian process, the background noise does not affect the bispectrum [34]. Single-frame spikes in the intensity trace with constant magnitude *I*_outlier_ and appearing with a constant rate *µ*_outlier_ (i.e., a Poisson process) will introduce frequency-independent offsets in both the power spectrum and the bispectrum (see Methods) [34]. To mitigate the impact of outliers, we fit both spectra with an additive constant offset term. For the power spectrum, the constant is absorbed in the constant due to the Gaussian background noise.

### Application to individual experimental intensity traces

We apply mCOAST to previously published experimental DNA-PAINT data of multiple DNA origami structures designed with different numbers of docking strands (*N*_max_ = 1, 4, 12, and 48) [26]. The DNA origamis were imaged at different imager strand concentrations (*c*_i_ = 5 nM, 10 nM, and 20 nM) with sufficiently low excitation power to ensure that blinking events are purely reflecting binding and unbinding of imagers [26, 36], not premature bleaching of the fluorophore before imager unbinding, and to reduce photoinduced loss of docking strands. The sample time is Δ*t* = 0.2 s in all experiments, and *T*_msr_ = 60 min for *c*_i_ = 5 nM, and *T*_msr_ = 30 min for *c*_i_ = 10 and 20 nM. Figure 1**e-h** shows the results of mCOAST for four different origami designs for the highest imager concentration *c*_i_ = 20 nM. The *N*_max_-values are upper bounds for the number of docking strands since imperfect strand incorporation into the origamis usually results in lower numbers, see, e.g., Fig. 1**i** for *N*_max_ = 4 where most origamis only contain 1 or 2 docking strands [26]. Origamis can, however, also misfold or aggregate, resulting in clusters with more binding sites than *N*_max_.

The precision of the estimates on *N* depends on the number of binding and unbinding events *n*_event_ during the course of the measurement, i.e., on the imager concentration *c*_i_. For the available experimental traces, reliable estimates for *N* are only obtained for the highest imager concentration (*c*_i_ = 20 nM) due to uncertainty on the estimates of the cumulants. The results for lower imager concentrations (*c*_i_ = 5 and 10 nM) are shown in Supplementary Note 1, Fig. S2. Here we also compared the mCOAST results with lbFCS for all imager concentrations [26]. lbFCS shows narrower distributions than mCOAST, especially for *N*_max_ = 48, which is expected as lbFCS depends on averaging the kinetics over all molecules in three experiments, while the mCOAST estimates are purely based on individual intensity traces without any prior knowledge. This is necessary when samples are heterogeneous, as in experiments in cellular contexts.

For all imager concentrations, mCOAST also provides estimates for the intensity unit *I*_on_, and the kinetic parameters 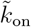 and *k*_off_ . As an example, Fig. 2**a-d** show histograms of 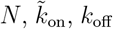, and *I*_on_, respectively, for *N*_max_ = 4 and *c*_i_ = 10 nM. All mCOAST estimates are performed independently on individual intensity traces, and all distributions show a single peak. The widths of the peaks are due to both the statistical uncertainty and the intrinsic variation of these parameters between different origami structures.

The results for 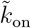 and *k*_off_ for all DNA origamis are summarized in Fig. 2**d-l**, showing the mean values of the parameters for different imager concentrations. As expected, scales linearly with imager concentration *c*_i_, with a slope of *k*_on_. The *k*_off_ -value is expected to be independent of imager concentration, which is indeed observed. For the mCOAST results, the fitted *k*_on_ values differ slightly between different origami designs, which is likely explained by docking strands on different positions on the structure experiencing different steric hindrance from the surroundings [37]. The kinetic rates are similar to those obtained in calibration experiments with lbFCS for *N* = 1 origami structures, i.e., *k*_on_ = 6.5 · 10^6^ M^−1^s^−1^ and *k*_off_ = 0.266 s^−1^ (dashed red lines in Fig. 2**e-l**) [26].

### Ensemble-averaging kinetic parameters for spots in the same field of view

As demonstrated above, mCOAST gives estimates for the number of targets and the kinetic parameter based on individual intensity traces without calibration experiments. However, if the estimates for the kinetic parameters 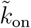 and *k*_off_ do not change between molecular clusters within the same sample or field of view, it can be assumed that the kinetic parameters are identical and their average values can be used when estimating the number of targets *N* . Inserting these averages in the analytical expressions for *I*_mean_ and *σ*^2^, the number of targets are

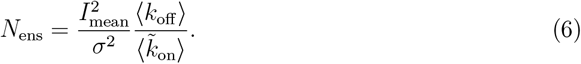

When estimating the number of targets in this way, *I*_mean_ and *σ*^2^ are from the individual clusters. Using the average values for the rates eliminates the need for the third-order cumulant, which is why C_3_ is not included in the expression.

As an example, Fig. 3**a-c** shows the estimated values for *I*_on_, *k*_on_, and *k*_off_ from individual DNA origamis across the entire field of view for *N*_max_ = 12 and *c*_i_ = 10 nM. The *I*_on_-values show a clear spatial variation, with maximum intensities located towards the upper left corner. This indicates a non-homogeneous Gaussian excitation intensity, typical for optical microscopy. In contrast, no systematic spatial variation is observed for the kinetic parameters 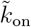 and *k*_off_ . Hence, we assume that all DNA origamis within the same field of view are imaged under identical experimental conditions, such as temperature and imager strand concentration. Consequently, we find the average values 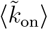 and ⟨*k*_off_⟩ and estimate the number of targets on individual DNA origamis using Eq. (6).

The results for *N*_ens_ for clusters within the same field of view are shown in Fig. 3 for imager concentrations *c*_i_ = 5, 10 and 20 nM. The overlapping peaks for different imager concentrations demonstrate the consistency of the method. For *c*_i_ = 20 nM, a comparison with Fig. 1**e–h** confirms that the calibrated kinetic parameter values increase precision. A comparison with lbFCS for all imager concentrations is shown in Supplementary Note 1, Fig. S4, showing good agreement between the methods. We emphasize that the lbFCS results depend on three measurements at different imager concentrations, while the mCOAST results are from a single imager concentration.

### Numerical simulations

We further investigate mCOAST by generating and analyzing simulated intensity traces for experimentally relevant parameters. Input parameters are therefore in all simulations *k*_on_ = 6.5 · 10^6^ M^−1^s^−1^, *k*_off_ = 0.266 s^−1^, *c*_i_ = 5 nM, 10 nM and 20 nM, 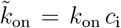, and Δ*t* = 0.2 s, identical to the values in [26] and similar to the values found in Fig. 2 with mCOAST. The procedure for simulating the state of a DNA origami, i.e., the number of bound imager strands versus time, is outlined in Supplementary Note 2.

First, we analyze the effect of background noise, e.g., from imager strands freely diffusing in the solution near the DNA origami. So, we simulate the state of a DNA origami with *N* ^(true)^ = 12 targets for an imager concentration *c*_i_ = 20 nM (Fig. 4**a,d**, black curves). The corresponding intensity traces *I*(*t*) are obtained by adding a zero-mean Gaussian white noise with standard deviation *σ*_bg_ to the simulated state of the origami (Fig. 4**a,d**, green curves). The background noise levels are *σ*_bg_ = 0.2*I*_on_ and *σ*_bg_ = *I*_on_, respectively. The corresponding histograms of the intensities *I*(*t*) for a measurement time *T*_msr_ = 60 min are shown in Fig.4**b**,**e**. For *σ*_bg_ = 0.2*I*_on_, distinct peaks are clearly visible in the intensity histogram. These peaks are absent for *σ*_bg_ = *I*_on_, where only a single, broad peak occurs.

Figure 4**c,e** show the estimated number of targets *N* based on individual traces [Eq. (1)] and on the average rates *N*_ens_ from all traces [Eq. (6)] for measurement times between 30 and 240 minutes. The background noise levels are *σ*_bg_ = 0.2*I*_on_ and *σ*_bg_ = *I*_on_, respectively. The means of the estimated values agree with the true number of targets for all measurement times, but, as expected, the spread in estimates is significantly larger for estimates based on individual traces [Eq. (1)] than for estimates with ensemble-averaged kinetic parameters [Eq. (6)]. For identical measurement times, the difference in estimates between the two noise levels is negligible. This demonstrates the ability of mCOAST to handle additive Gaussian white noise. The estimates from individual traces [Eq. (1)] for *N* ^(true)^ = 4, 12, and 20 for *σ*_bg_ = *I*_on_ are shown in Supplementary Note 1, Fig. S5 for different measurement times and imager concentrations. The distribution of estimates is for all cases centered around the input value, with the standard deviation decreasing as imager concentration and measurement duration increase. Accordingly, higher imager concentrations and longer time traces result in more precise quantification of the number of targets, *N* .

In addition to Gaussian background noise, peaks in intensity traces can occur, e.g., due to unspecific binding of imager strands to the DNA origami, or other objects passing close by the cluster. We model the outliers in the intensity trace by adding spikes to the intensity trace with a fixed rate *µ*_outlier_ (a Poisson process) and intensity *I*_outlier_. Figure 4**g-i** show the results of simulations for *N* ^(true)^ = 12, *σ*_bg_ = *I*_on_, *T*_msr_ = 60 min, *µ*_outlier_ = 0.03 s^−1^, and *I*_outlier_ = 4*I*_on_. The red curve in Fig. 4**g** shows the occurrences of outliers, which are not visible in the intensity histogram (Fig. 4**g**). In Fig. 4**h**, the number of targets are estimated with values for *σ*^2^ and C_3_ from fits to the power spectra and the bispectra, but also from direct calculations of the moments with 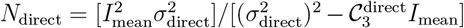 (see the section ‘Cumulants from experimental intensity traces’). The *N*_direct_-estimates consistently overestimate the number of targets, as outliers in the intensity trace can significantly influence the value of direct calculations, especially the third-order cumulant [33]. The effect of outliers can be absorbed in the fits of the power and bispectra. This stresses the importance of obtaining the cumulants from fits to the power and, in particular, the bispectrum, and not from direct calculations.

The numerical simulations show that mCOAST correctly estimates the number of targets, even when no distinct peaks appear in the intensity histogram (as required for lbFCS+ [27]) due to background noise, and with outliers occurring at a fixed rate (Fig. 4). The precision of mCOAST increases with imager concentration, resulting in narrower distributions for the estimates of *N*, as opposed to, e.g., qPAINT that starts underestimating *N* for higher concentrations due to overlapping events. Tuning the DNA hybridization kinetic parameters 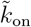 and *k*_off_ is technically also an option, but these are usually set by the chosen DNA sequences and the imaging buffer. Finally, we note that numerical simulations can be used to estimate uncertainties in the fitted parameters by using the fitted values as input and performing subsequent fitting.

## Discussion

mCOAST extends previous time series-based methods for molecular counting [22, 26, 27]. qPAINT requires that 1) the rate 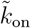 is known from a calibration experiment or from the literature, 2) the imager concentration is sufficiently low so that binding events do not overlap in time, and 3) low noise levels so the duration of time intervals with no binding events can be clearly identified [22]. lbFCS also relies on a known value for *k*_on_, and the number of targets in a cluster is determined as 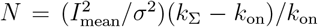 [26]. This is similar to Eq. (6), but with both *σ*^2^ and *k*_Σ_ obtained from a fit to the autocorrelation function. As lbFCS does not require clearly identifiable dark times in the trajectory, it is less sensitive to noise than qPAINT and works for higher imager concentrations. lbFCS+, a calibration-free method, relies on scaling of the peaks in the intensity histogram, so the standard deviation of the background noise must be less than the intensity of individual binding events (see, e.g., Fig. 4**b,d**), and it is only demonstrated for *N* < 10. Blinx, a recent calibration-free alternative based on a Bayesian model [38], fits the individual steps in the intensity trace, and extracts the number of targets and the kinetic parameters. However, blinx requires very low noise levels to avoid overfitting the number of targets and that *k*_on_ > *k*_off_, i.e., the opposite regime of most experiments, e.g., qPAINT and lbFCS [22, 26].

The main advantages of mCOAST compared to existing approaches are that mCOAST works on individual clusters of localizations, does not require any prior knowledge regarding the binding kinetics, gets more precise with increasing imager concentration, does not require fine-tuning of the imager concentration, and can handle background noise and outliers. The data analysis is fast and transparent, as parameter estimation relies only on the first three cumulants and the correlation time 1*/k*_Σ_, all obtained from fits to the cumulant spectra of the intensity trace from a single cluster of targets.

The trade-off for the lack of calibration is trajectories with more binding-unbinding events per target compared to qPAINT and lbFCS to achieve similar precision. This can be obtained by longer acquisition times, higher imager concentrations, or modifications of the DNA strands. Utilizing additional clusters within the same field of view enables the application of mCOAST at lower imager strand concentrations or with shorter measurement times, but loses the ability to detect kinetic heterogeneity.

The use of the power spectrum to extract *σ*^2^ and *k*_Σ_ has certain advantages over the autocorrelation [26],[27]. For a time series of finite length, the values in the autocorrelation function are inherently correlated, and a fit to the autocorrelation function requires a carefully chosen maximum time lag. In contrast, the power spectral values are asymptotically independent and exponentially distributed around their expected values [35]. Hence, the parameters can be estimated by fitting the power spectrum with maximum likelihood estimation and do not depend on a user-defined maximum time lag.

Our theoretical framework (see Supplementary Note 2) can be further extended to account for effects such as time-averaging of the signal, which arises from the finite integration time of the camera. This can be achieved by incorporating temporal filters into the analysis of both the power spectrum and bispectrum [34, 39]. This becomes relevant if the integration time significantly exceeds the characteristic time 1*/k*_Σ_, which is the opposite regime of what is considered here.

In conclusion, mCOAST enables the necessary next step to bring molecular counting to challenging biological samples where the binding kinetics are unknown and cannot be calibrated, or the molecular clusters are not fixated. Here, parameter estimation must be at the single-cluster level due to high diversity among molecular clusters, and tuning of the imager concentration is challenging. Beyond its application for DNA-PAINT, mCOAST is applicable to all molecular two-state interactions fulfilling the assumption of independent and identical targets, and constant kinetic parameters, e.g., PAINT with proteins, peptides, enzymes, or antibodies [40].

## Methods

### Extraction of intensity traces from raw image stacks

The description of the DNA-PAINT experiments can be found in [26], where the (*x, y*)-positions of the DNA origamis of interest from a stack of raw images were found using the software Picasso [41]. For the specific parameters for the selection of clusters, see Suppl. Table 3 in [26]. In our analysis, we use the localization clusters analyzed and filtered in [26].

In the present paper, we get the intensity trace per cluster by integrating the 3 × 3 pixel intensities centered around the (*x, y*)-coordinate of the cluster. We obtain the background intensity in every frame by taking the median of the 7 × 7 border pixels around each cluster. By subtracting the background intensity in each frame, we obtain the intensity trace per cluster. In contrast to the traces obtained with Picasso, where the recorded intensity in each frame results from a Gaussian fit of the spot, which may suffer from spot detection errors and fitting uncertainty (see Supplementary Note 1, Fig. S1**f**). Background subtraction, a common and trivial task for this type of experiment, constitutes the only data processing in mCOAST. We finally exclude clusters less than five pixels apart to ensure that the signals from neighboring clusters do not affect the analysis.

### Recorded intensities and the discrete Fourier transform

The spot intensities in each frame compose the intensity trace *I*_1_, *I*_2_, *I*_3_ …, *I*_*n*_, where *I*_*j*_ = *I*(*t*_*j*_) is the background-corrected intensity of frame *i* recorded at time *t*_*j*_ = *j*Δ*t*. Here Δ*t* is the sample time, *T*_msr_ = *n*Δ*t* is the total measurement time, and *n* is the number of samples in the trace.

The discrete Fourier transform of the intensity trace is defined as 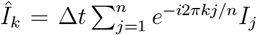 for *k* = 0, 1, …, *n* − 1, and the frequency as *f*_*k*_ = *k/T*_msr_.

### Fit of power spectrum

The discrete, experimental power spectrum is defined as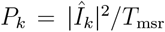. The experimental power spectrum is fitted to the expected value of the theoretical power spectrum (see Supplementary Note 2),

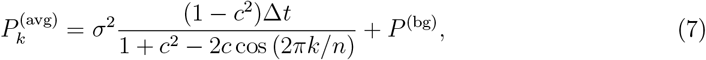

with 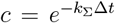. The first term is the contribution from the targets. The last, frequency-independent term 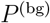 is the total contribution from the Gaussian background, 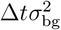, and outliers ([34], Eq. 3.25b).

In order to fit the power spectrum using maximum likelihood, we use that under not too strict conditions, hold that the discrete power spectral values *P*_*k*_ are asymptotically independent and exponentially distributed with scale parameter equal to the average value 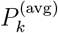 [35], i.e., the probability to get the value *P*_*k*_ is Prob 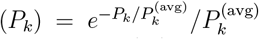. The likelihood function is defined as 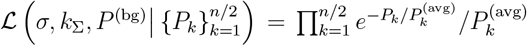 . According to the Maximum Likelihood principle, the best estimates for the parameters are those that maximize the likelihood, or equivalently, minimize the negative log-likelihood function. We only fit frequencies up to the Nyquist frequency *f*_Nyq_ = *n/*(2Δ*t*) due to the periodicity of the power spectrum [33].

The exponential distribution of the power spectral values around their average values can serve as a diagnostic criterion for evaluating the quality of the fits, as 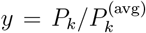 is expected to be exponentially distributed, Prob(*y*) = *e*^−*y*^. Supplementary Note 1, Figure S6-9 shows the analysis for individual, simulated trajectories for different numbers of targets *N* ^(true)^ and different noise levels *σ*_bg_ for parameters corresponding to *c*_i_ = 20 nM. Panels **d** show that 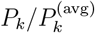 are indeed exponentially distributed.

### Fit of bispectrum

We calculate the experimental bispectrum following section IVB in [34]. In brief, the intensity trace *I*_*j*_ (*j* = 1, 2, 3, … *n*) is divided into *K* segments, each having *m* = *n/K* time points. We set *K* = 20 for all analyzed traces, but the results do not depend critically on this parameter. We then subtract the mean value from each segment and calculate its discrete Fourier transform. For a time-translational invariant system, the discrete bispectrum is 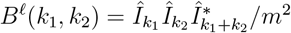, where *ℓ* = 1, 2, … *K, k*_1,2_ = 0, 1, …, *m/*2, and […]^∗^ the complex conjugate ([34], Eq. 4.12b). The final experimental bispectrum is the average over the *K* bispectra,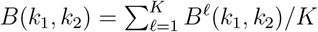.

The average value of the bispectrum for the two-state jump process is (see Supplementary Note 2)

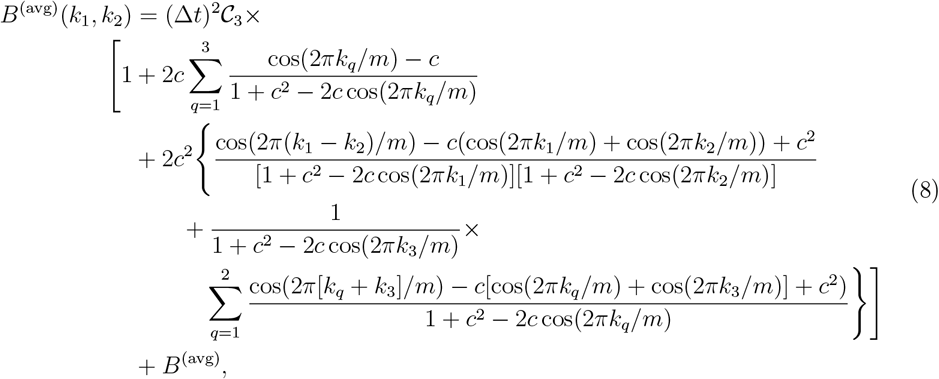

where *k*_3_ = *k*_1_ + *k*_2_ and *B*^(avg)^ is a frequency-independent constant due to delta-shaped intensity outliers with constant magnitude that occur with a fixed rate in the time trace [34]. Notice that the bispectral values *B*(*k*_1_, *k*_2_) are complex valued, but the expected value of the imaginary part is zero for all frequencies. So, we only fit the real part of the spectrum.

The estimated values for the bispectrum *B*(*k*_1_, *k*_2_) tend to be normally distributed for sufficiently large *m* and *K* (Central Limit Theorem). The mean value is approximately the average value of the bispectrum, and the variance is [34, 42]

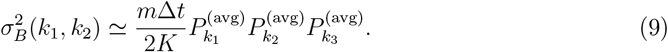

Here *P* ^(avg)^ is defined in Eq. (7), but with *n* replaced with *m*. We fit the real part of the bispectrum to Eq. (8) using a weighted least-squares algorithm with weight factors 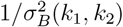. Due to the symmetries of the bispectrum, we only consider the triangular region 0 ≤ *k*_2_ ≤ *k*_1_, *k*_1_ + *k*_2_ ≤ *m/*2 [34]. The fit parameters are the third-order cumulant 𝒞_3_ and *k*_Σ_ (recall that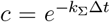). The *k*_Σ_ is determined with higher precision from the power spectrum fit, so we adopt the value obtained from the power spectrum when estimating the model parameters. The expected normal distribution of the power spectral values around their expected values can serve as a diagnostic criterion for evaluating the quality of fits. That is, 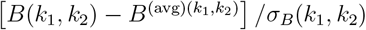 is expected to be normally distributed with zero mean and unit standard deviation. Supplementary Note 1, Figure S6-9 shows the analysis for individual, simulated trajectories for different numbers of targets *N* ^(true)^ and different noise levels *σ*_bg_ for parameters corresponding to *c*_i_ = 20 nM. Panels **f** show that 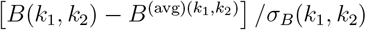 indeed follows a standard normal distribution.

## Data availability

The data that support the findings of this study are available from the corresponding authors upon reasonable request.

## Code availability

The code used in this study is available on Github: github.com/TeunHuijben/mcoast.

## Acknowledgements

The authors thank Petra Schwille, Ralf Jungmann, Florian Stehr, and Johannes Stein for kindly sharing the raw origami data from [26]. This project has received funding from the European Union’s Horizon 2020 research and innovation program under the Marie Skłodowska-Curie grant agreement SuperCol (grant agreement no. 860914), and from the Novo Nordisk Foundation under the New Exploratory Research and Discovery program (grant agreement no. NNF21OC0068622).

## Author contributions

T.A.P.M.H. and J.N.P. conceptualised the idea, derived theory, ran simulations and performed data analysis. T.A.P.M.H. made all figures and wrote the initial version of the manuscript. J.N.P. and R.M. supervised the work. All authors commented on the manuscript.

## Supplementary Notes

### 1 Supplementary Note 1: Figures and Tables

**Supplementary Figure 1.**
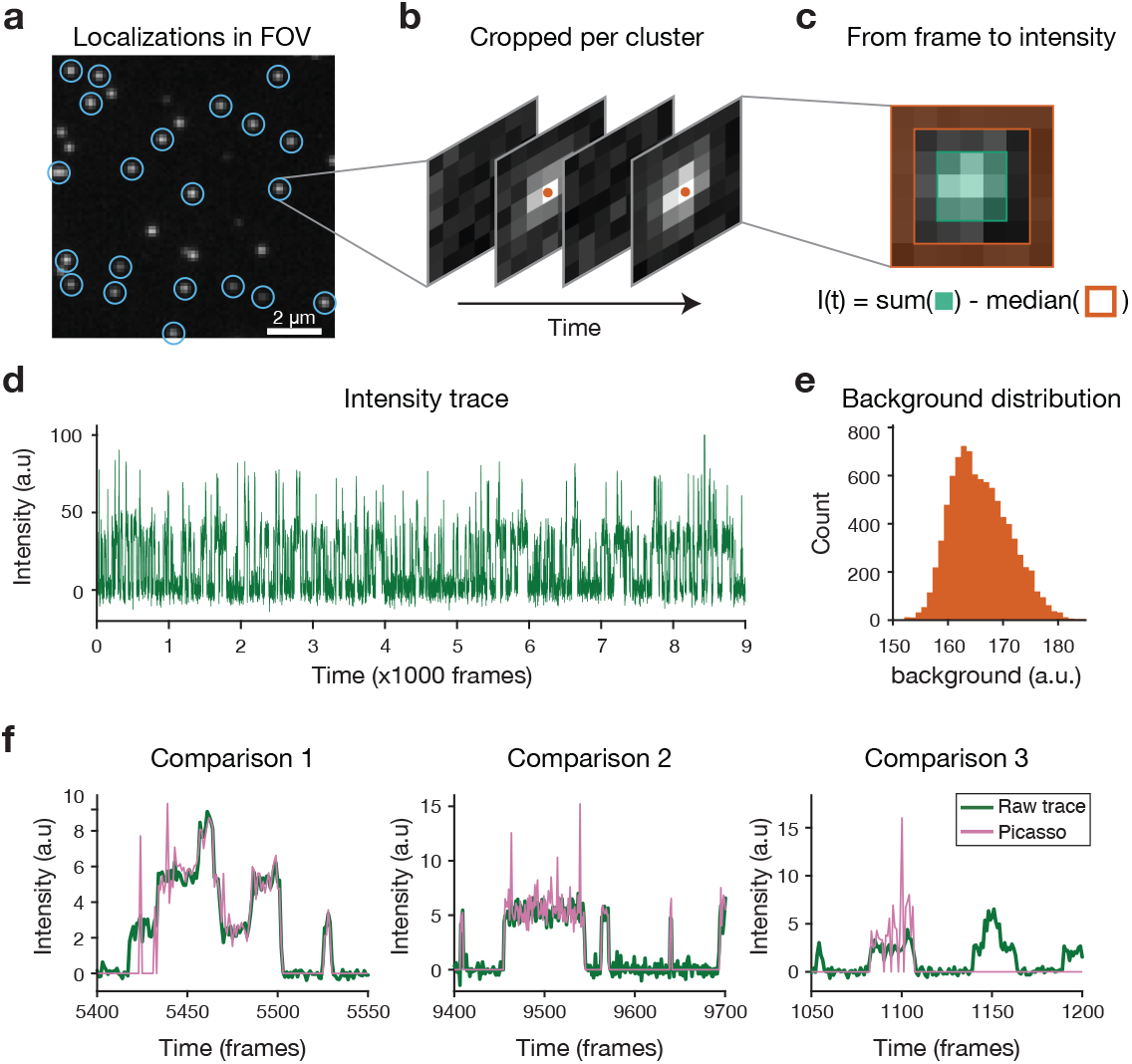
Extraction of raw intensity traces. **a** To extract the intensity traces from a stack of raw images, the software Picasso ran through the stack of localizations to find the (*x, y*)-positions of the spots in every frame. After grouping the localizations per origami structure and picking the groups of interest, the time-averaged (*x, y*)-position of the clusters was calculated. Applying drift correction gives the (*x, y*)-position of clusters at every frame (blue circles). **b** For every cluster, we crop the 7 × 7 pixels around the drift-corrected position in all frames. Picasso would run a spot detection algorithm, and fit a 2D Gaussian to every frame where a spot is detected, to obtain the position (red dots) and intensity of every event, which can be rendered as an intensity time-trace. **c** However, a more robust way to extract intensity traces is by directly integrating the pixel intensities in the cropped images. Per frame, we calculate the background-subtracted intensity value by summing the inner 3 × 3 pixels (green shaded area) and subtracting the median of the 7 × 7 border pixels (red shaded area). **d** This procedure is repeated for every frame, resulting in the intensity time trace from a single cluster of localizations. **e** Histogram of the background intensities from a single trace. **f** Example section of three intensity time traces, either obtained by our method as explained here (‘raw trace’) or based on the fitted localizations as done in Picasso. The examples clearly highlight the robustness of the raw intensity traces, since the Picasso traces either have outliers with high intensities due to erroneous fitting, or miss events due to suboptimal spot detection algorithms.

**Supplementary Figure 2.**
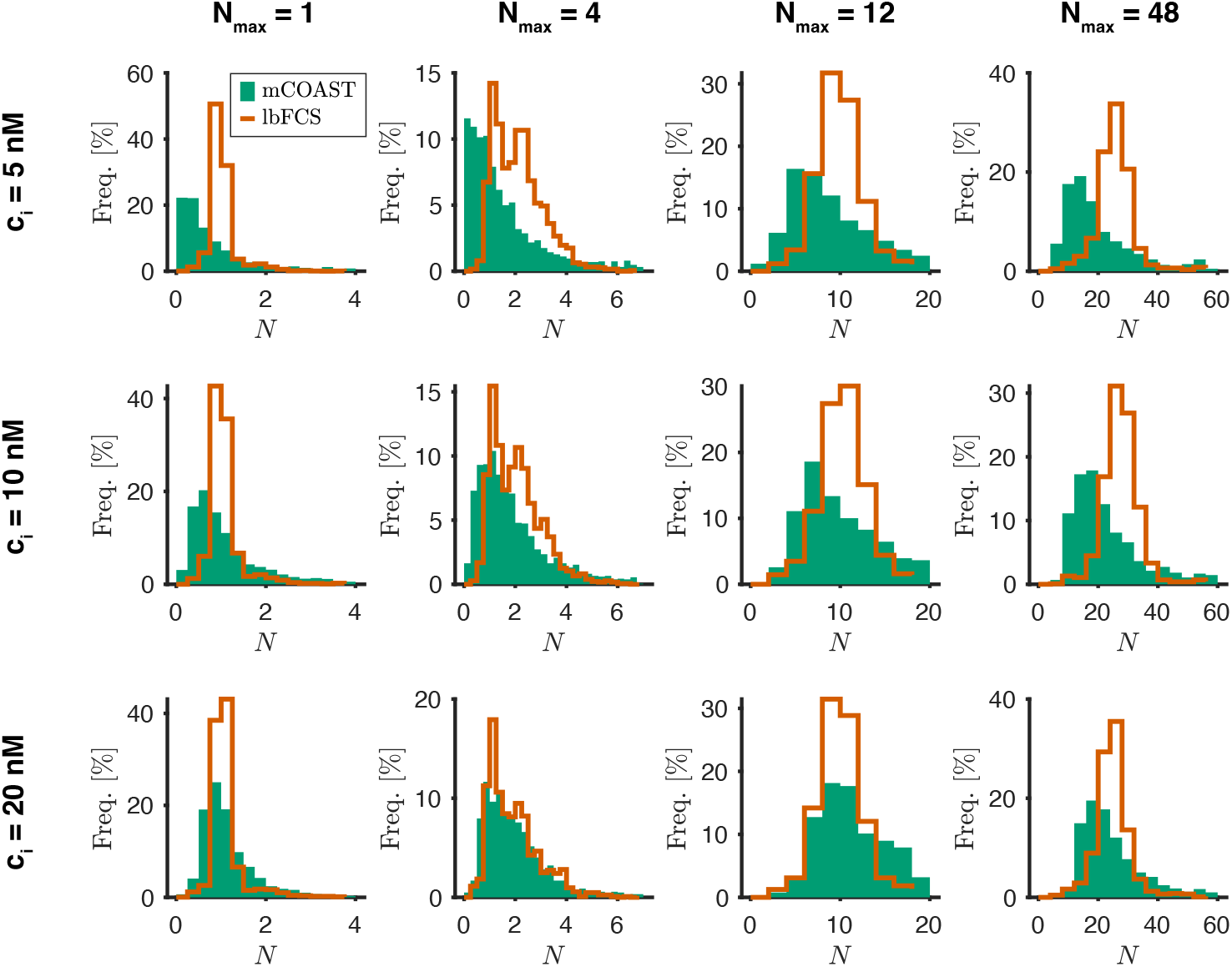
mCOAST estimates for low imager concentrations. Same as Fig. 1**e-h**, but including imager concetrations *c*_i_ = 5 nM (top row) and *c*_i_ = 10 nM (bottom row). Results for lbFCS (red curve) are shown for comparison [1]. The number of traces are stated in Supplementary Note 1, Table S1.

**Supplementary Figure 3.**
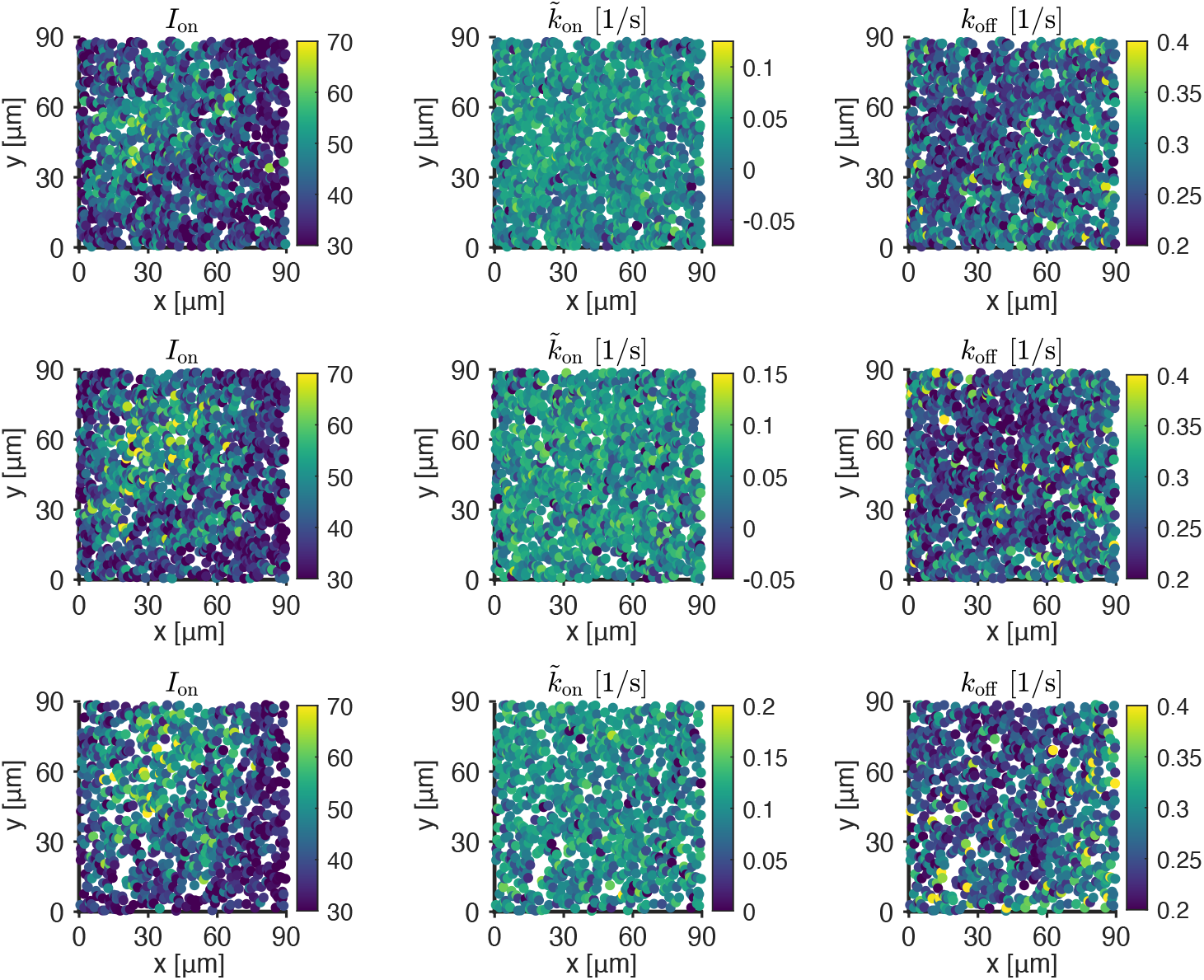
Estimated intensity of single binding events over the field of view. Scatter plot showing every analyzed cluster of localizations in the entire field of view as a single dot. The expected number of targets is *N*_max_ = 12, and the imager concentrations are *c*_i_ = 5 nM (top row), *c*_i_ = 10 nM (middle row), and *c*_i_ = 20 nM (bottom row).

**Supplementary Figure 4.**
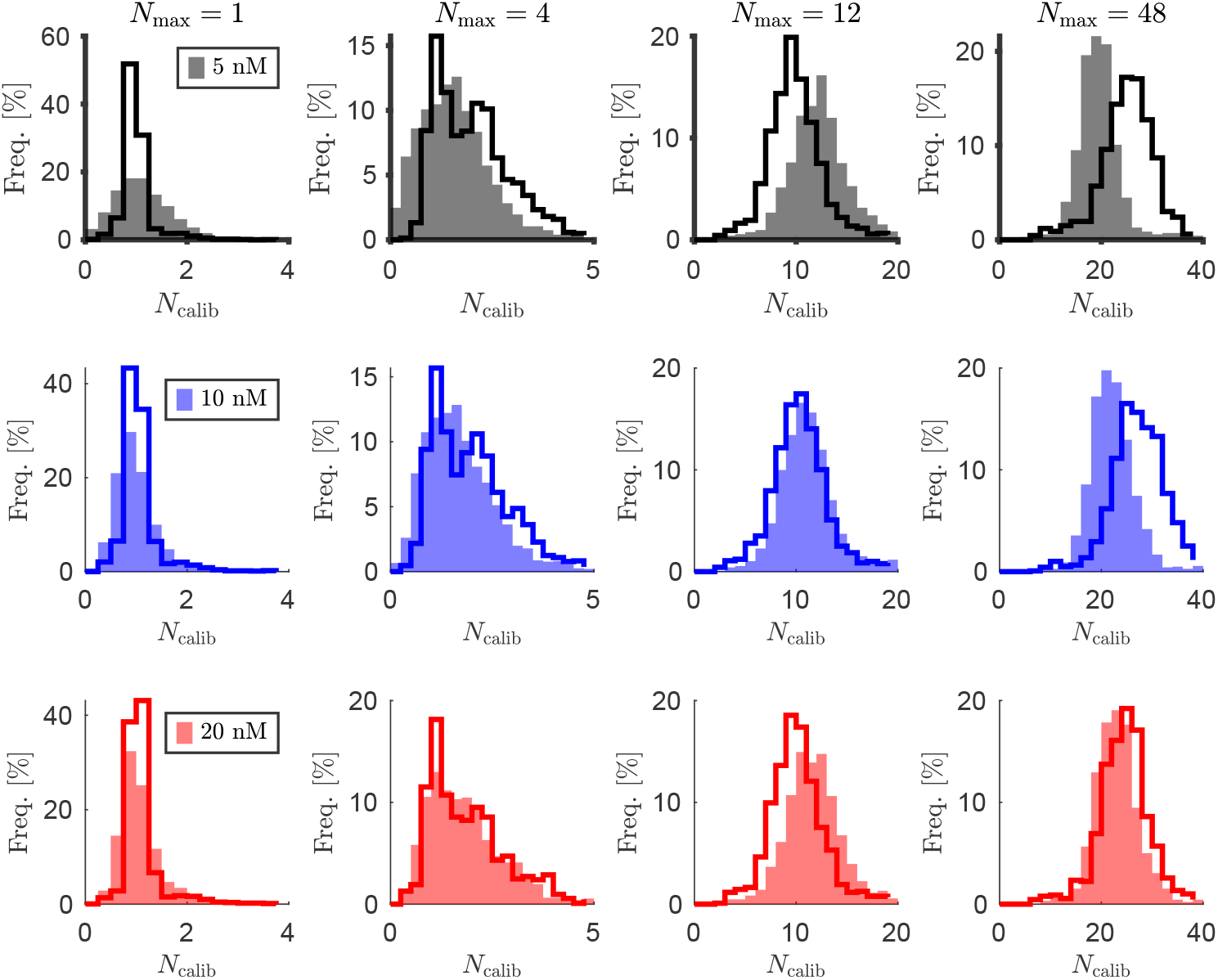
Ensemble-averaged mCOAST estimates versus lbFCS. Comparison of results with mCOAST for ensemble-averaged values for 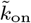 and *k*_off_ [Eq. (6) in the main text, shaded area] and lbFCS (full lines) for the imager concentrations *c*_i_ = 5, 10, and 20 nM (upper to lower row), and *N*_max_ = 1, 4, 12, and 48 (left to right column). The number of traces is stated in Supplementary Note 1, Table S1.

**Supplementary Figure 5.**
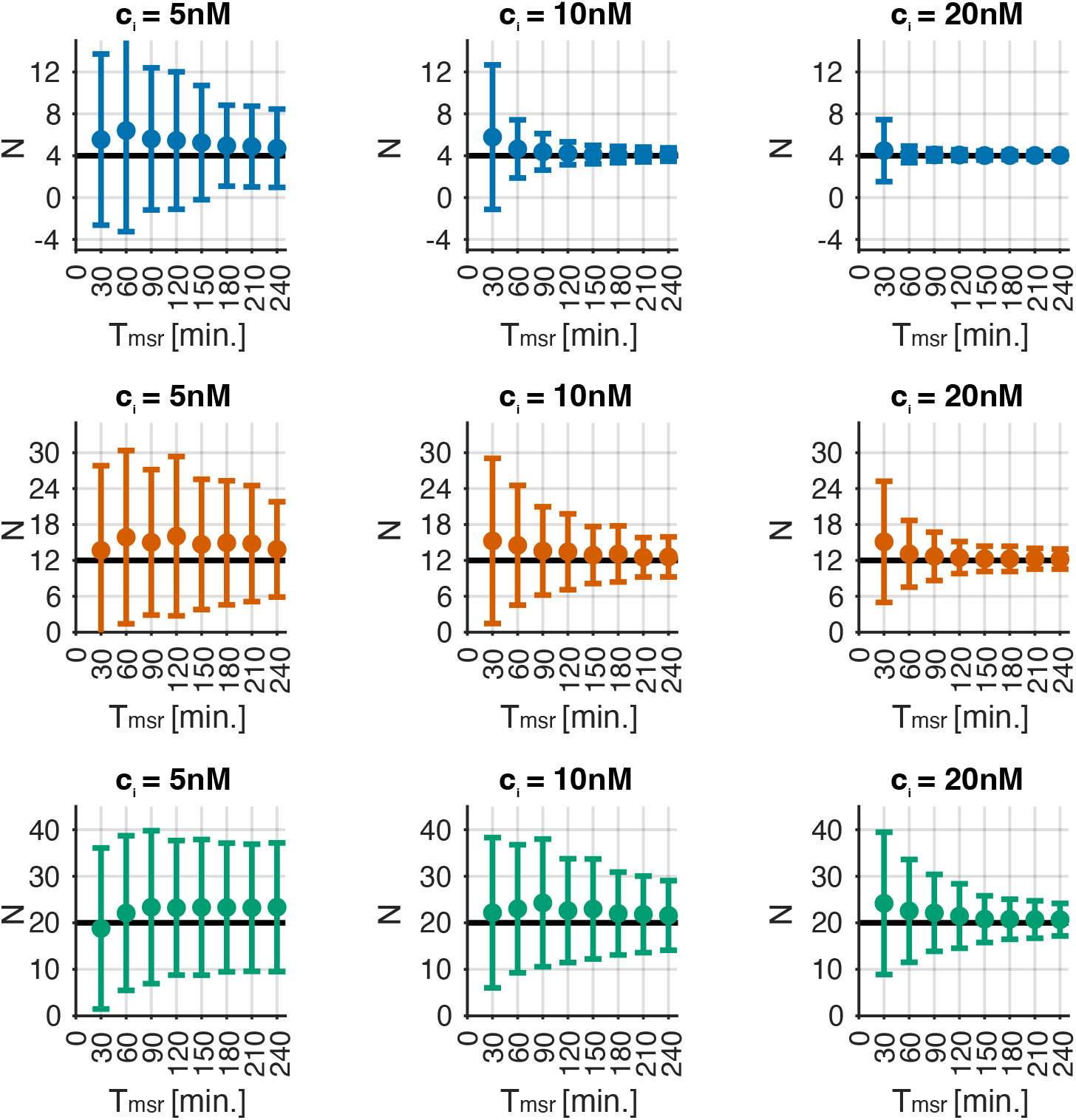
Uncertainties on estimated number of targets for different imager concentrations. Estimated number of targets *N* versus measurement time *T*_msr_ based on individual traces [Eq. (1) in the main text]. The true values of the number of targets are *N* ^(true)^ = 4 (top row), *N* ^(true)^ = 12 (middle row), and *N* ^(true)^ = 20 (bottom row). Other input parameters are *k*_on_ = 6.5 · 10^6^ M^*−*1^s^*−*1^, *k*_off_ = 0.266 s^*−*1^, *c*_i_ = 5 nM, 10 nM and 20 nM, 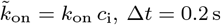, and *σ*_bg_ = *I*_on_. Simulations are repeated 1000 times for all parameter sets. Full dots are means, and error bars are standard deviations.

**Supplementary Figure 6.**
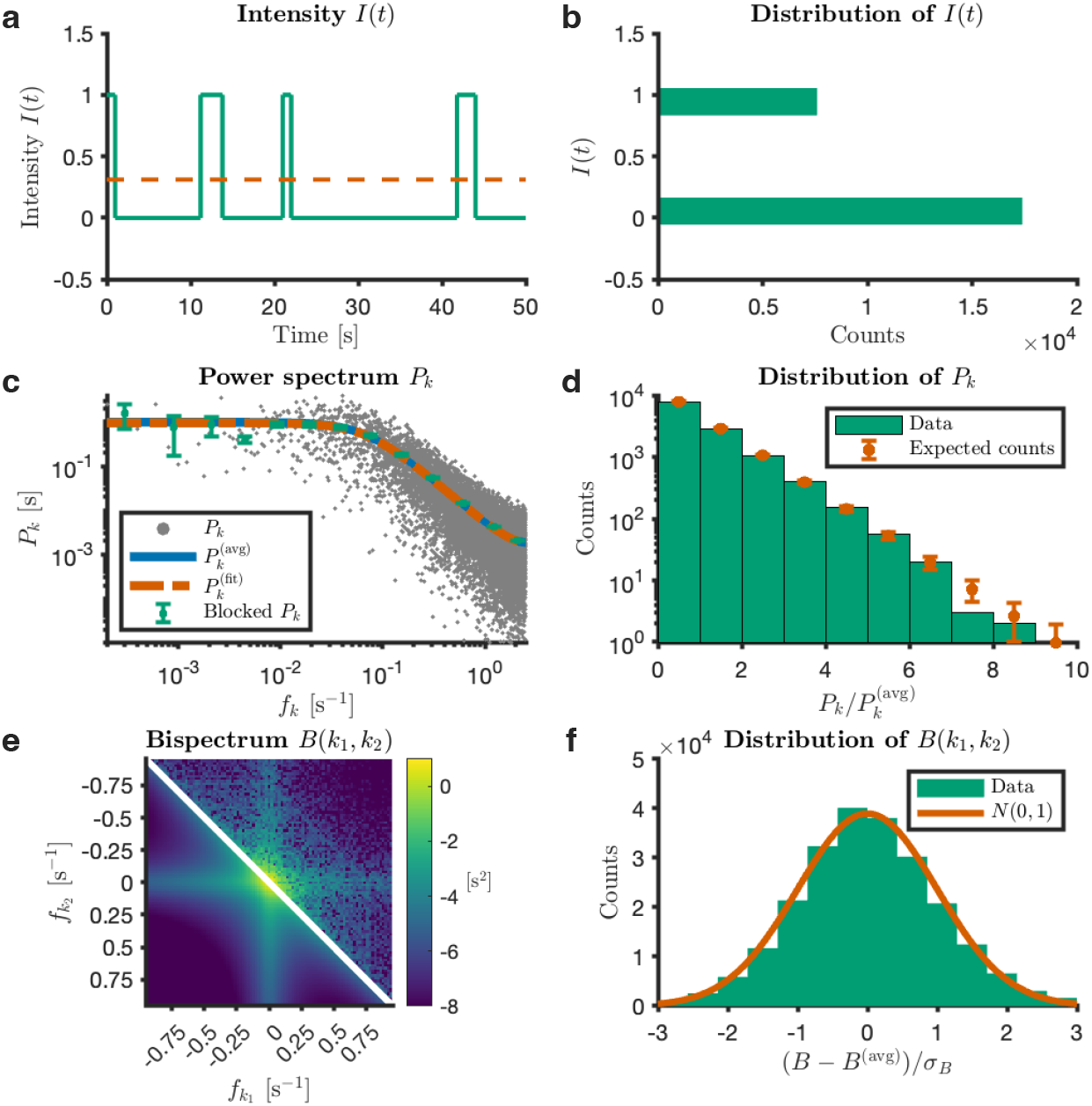
Simulated intensity trace without noise for a single target. Simulation of the intensity *I*(*t*) for the parameters *N* ^(true)^ = 1, *k*_on_ = 6.8 ·10^6^ M^*−*1^s^*−*1^, *k*_off_ = 0.3 s^*−*1^, *c*_i_ = 20 nM (resulting in 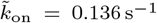), Δ*t* = 0.2 s, *I*_on_ = 1, and *T*_msr_ = 5000 s. **a** First 50 s of the intensity *I*(*t*) (green), with the red dotted line representing the theoretical mean. **b** Histogram of intensities of *I*(*t*). **c** Experimental power spectral values *P*_*k*_ [grey dots, Eq. (S20)], the block-averaged experimental power spectral values with the standard error on the mean as error bars (green), the theoretical power spectral values 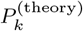 (blue), and the fitted power spectrum 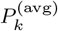 (red) from Maximum Likelihood estimation of the parameters (see Methods). **d** Histogram of the ratios 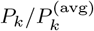. The red dot marks the theoretically expected number of counts in each bin. Error bars are the standard deviation (square roots of the expected number of counts). **e** Simulated bispectrum *B*(*k*_1_, *k*_2_) (upper half), and fit to *B*^(avg)^(*k*_1_, *k*_2_) from Eq. (8) in the main text (lower half). **e** Scaled residuals 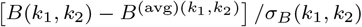 (see Methods). The red curve shows a standard normal distribution. The mCOAST estimates for this trace are 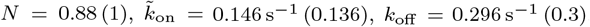, and *I*_on_ = 1.04 (1). The values in the parentheses are the input values in the simulation.

**Supplementary Figure 7.**
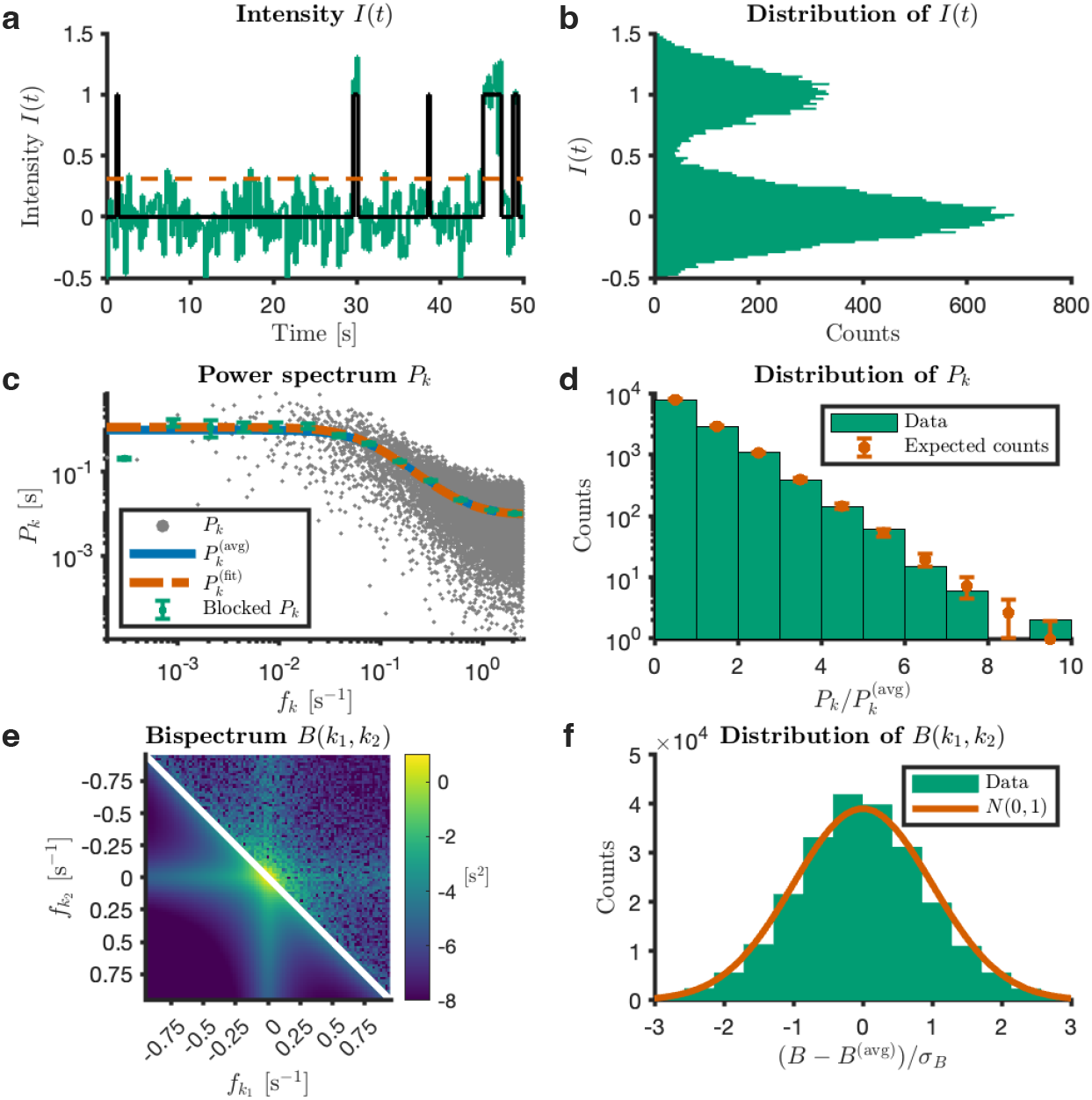
Simulated intensity trace with noise for a single target. Same parameters as in Fig. 6, but with an additional Gaussian background intensity *I*^bg^ with zero mean and standard deviation *σ*_bg_ = 0.2 *I*_on_. The mCOAST estimates of this trace are 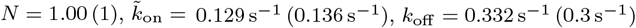, and *I*_on_ = 1.00 (1). The values in the parentheses are the input values in the simulation.

**Supplementary Figure 8.**
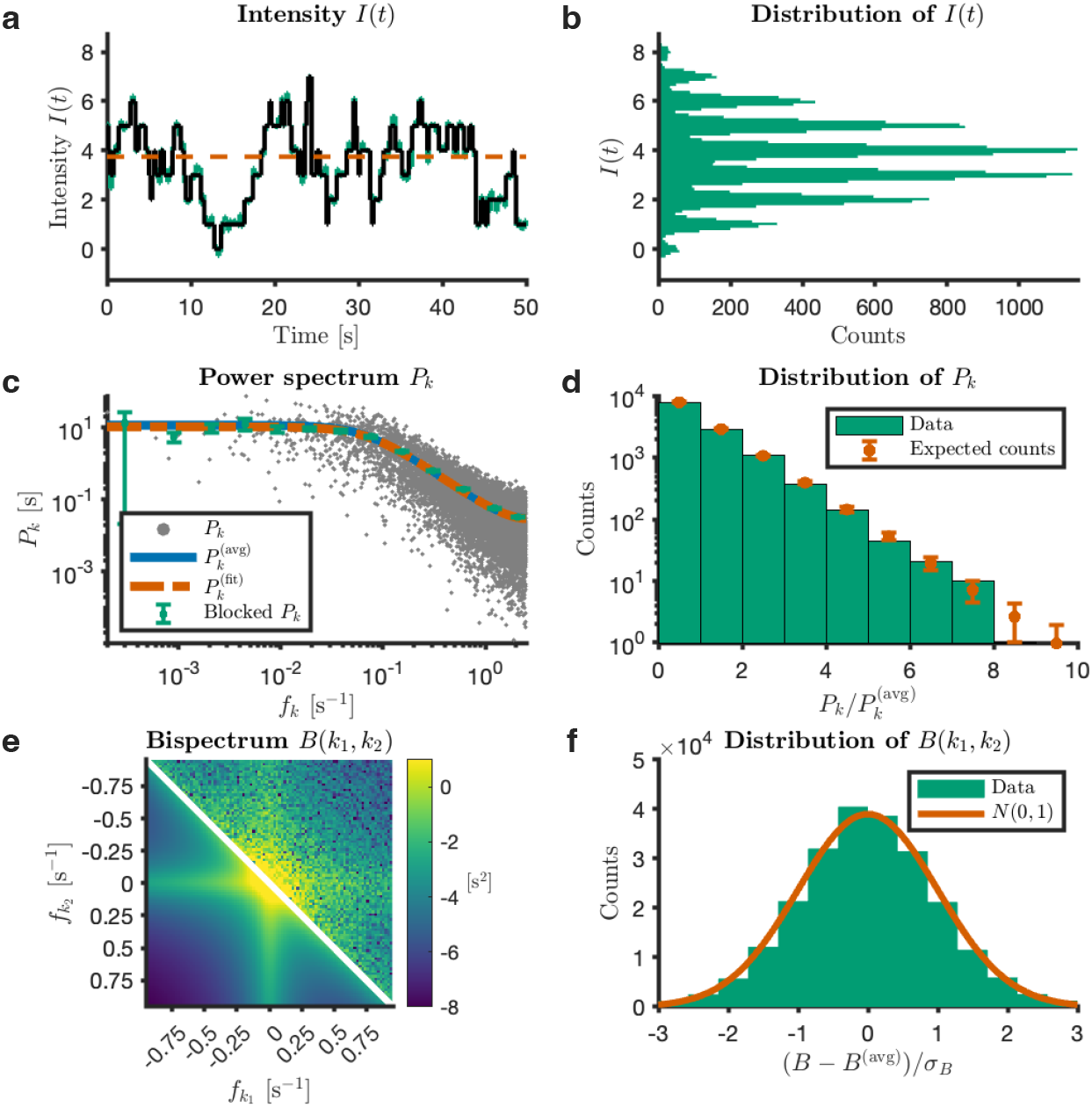
Simulated intensity trace with low noise for *N* = 12 targets. Same parameters as in Fig. 6, except *N* ^(true)^ = 12 and an added Gaussian background noise with zero mean and standard deviation *σ*_bg_ = 0.2 *I*_on_. The mCOAST estimates of this trace are 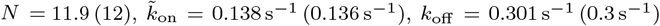, and *I*_on_ = 0.99 (1). The values in the parentheses are the input values in the simulation.

**Supplementary Figure 9.**
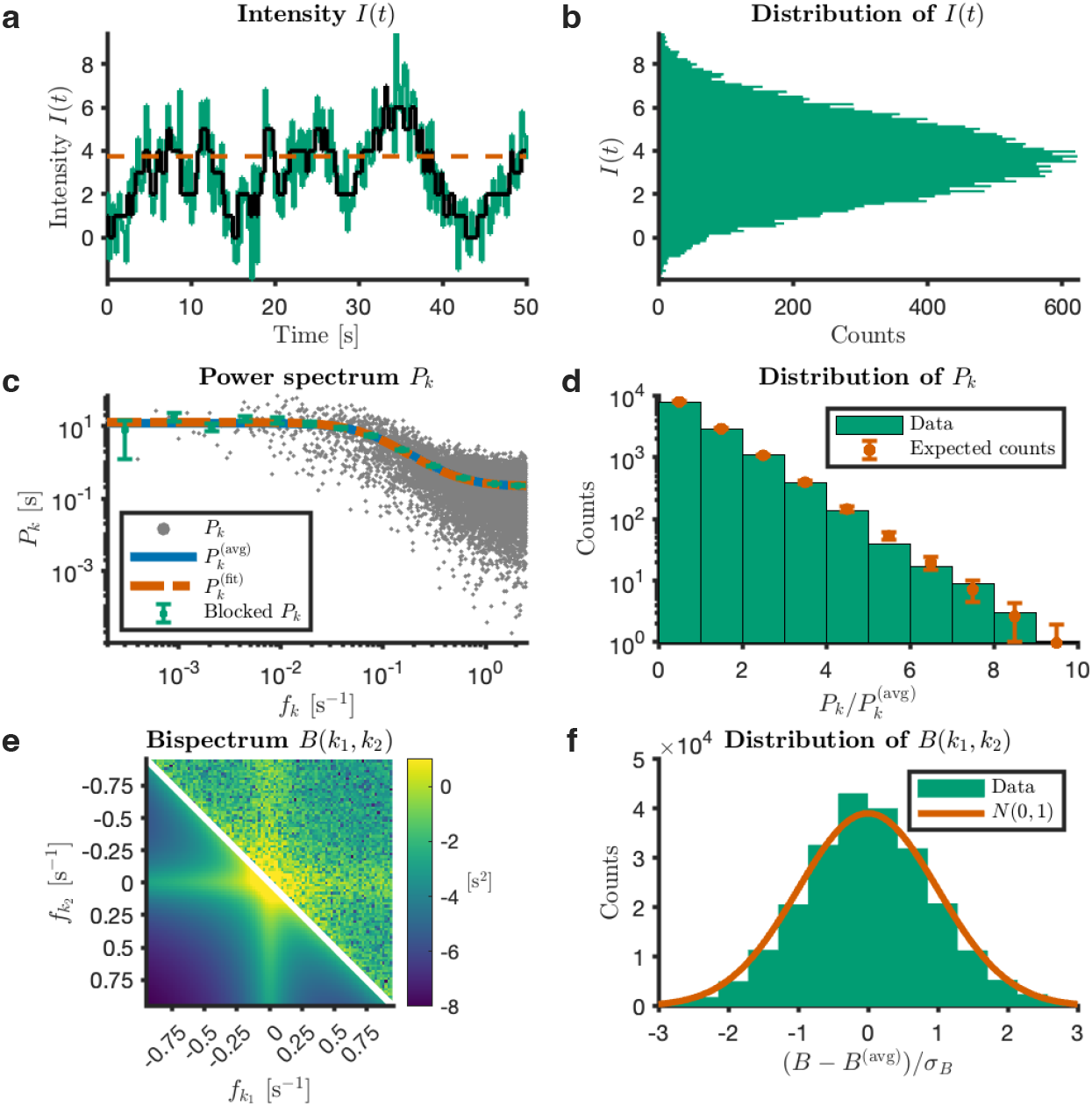
Simulated intensity trace with high noise for *N* = 12 targets. Same parameters as in Fig. 8, except *σ*_bg_ = *I*_on_. The mCOAST estimates of this trace are 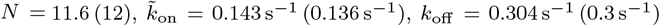, and *I*_on_ = 1.01 (1). The values in the parentheses are the input values in the simulation.

**Supplementary Table 1:** Total no. of clusters more than 5 pixels apart, no. of traces in Fig. 3 (mCOAST), and in Figs. S2 and S4 (mCOAST and lbFCS).

| $N_{\max}$ | $c_i$ | Total no. of traces | mCOAST | lbFCS |
| --- | --- | --- | --- | --- |
| 1 | 5 | 2166 | 2155 | 2157 |
| 1 | 10 | 2305 | 2289 | 2289 |
| 1 | 20 | 2297 | 2288 | 2285 |
| 4 | 5 | 2303 | 2294 | 2276 |
| 4 | 10 | 1956 | 1946 | 1933 |
| 4 | 20 | 1271 | 1255 | 1251 |
| 12 | 5 | 1461 | 1349 | 1413 |
| 12 | 10 | 1064 | 1363 | 1397 |
| 12 | 20 | 694 | 999 | 1035 |
| 48 | 5 | 1436 | 678 | 667 |
| 48 | 10 | 1436 | 1398 | 1393 |
| 48 | 20 | 919 | 885 | 895 |

**Supplementary Table 2:**
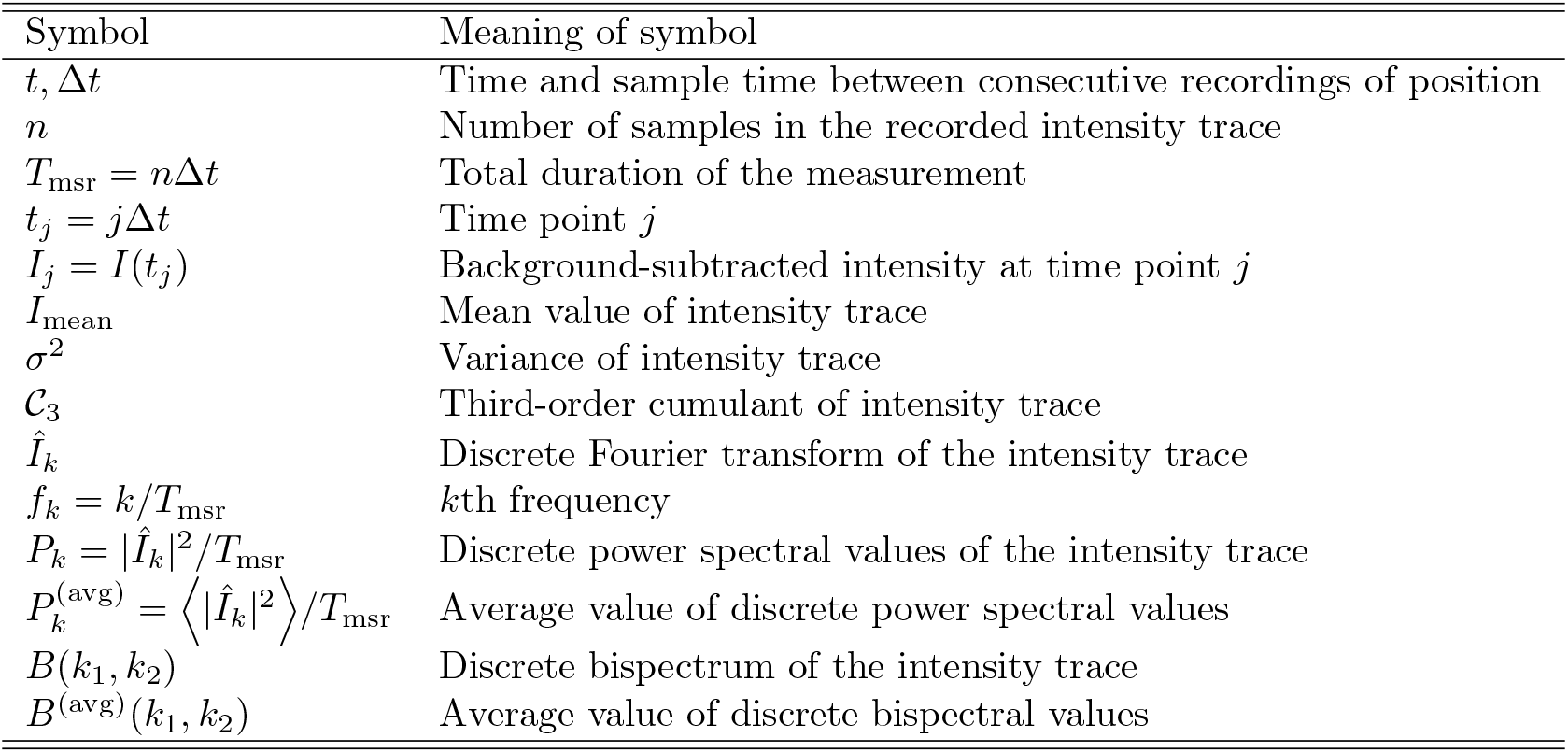
Notation pertaining to the recorded intensity trace.

**Supplementary Table 3:** Notation pertaining to the state function *S*(*t*)

| Symbol | Meaning of symbol |
| --- | --- |
| $S(t), S_j = S(t_j)$ | State function $S$ at time $t$ and $t_j$, respectively |
| $k_{\text{on}}$ | Molar association rate |
| $c_i$ | Imager concentration |
| $\tilde{k}_{\text{on}} = c_i k_{\text{on}}$ | Rate for the 0 to 1 transition (effective association rate) |
| $k_{\text{off}}$ | Rate for the 1 to 0 transition (dissociation rate) |
| $k_{\Sigma} = \tilde{k}_{\text{on}} + k_{\text{off}}$ | Sum of rates |
| $k_{\Delta} = \tilde{k}_{\text{on}} - k_{\text{off}}$ | Difference between rates |
| $P_q(t)$ | Probability of state $q$ at time $t$ ($q = 0, 1$) |
| $P_{q \rightarrow q'}(t)$ | Probability of state $q$ at time 0 and $q'$ at time $t$ ($q, q' = 0, 1$) |
| $p_q$ | Probability of state $q = 0, 1$. |
| $c = e^{-k_{\Sigma} \Delta t}$ | Scale factor |
| $S_{\text{mean}}$ | Mean value of $S(t)$ |
| $\sigma_S^2$ | Variance of $S(t)$ |
| $\mathcal{C}_3^{(S)}$ | Third-order cumulant of $S(t)$ |
| $\phi_2(t, t')$ | Two-point correlation function |
| $\phi_3(t, t', t'')$ | Three-point correlation function |
| $C_2(t, t')$ | Second-order cumulant function (auto-covariance function) |
| $C_3(t, t', t'')$ | Third-order cumulant function |
| $P^{(S, \text{avg})}(\omega)$ | Continuous power spectrum of $S(t)$ (Fourier transf. of $C_2$) |
| $P_k^{(S, \text{avg})}$ | Discrete power spectrum of $S_j$ (Fourier transf. of $C_2$) |
| $B^{(S, \text{avg})}(\omega_1, \omega_2)$ | Continuous bispectrum of $S(t)$ (Fourier transf. of $C_3$) |
| $B^{(S, \text{avg})}(k_1, k_2)$ | Discrete bispectrum of $S_j$ (Fourier transf. of $C_3$) |

### 2 Supplementary Note 2: Mathematical Derivations

#### 2.1 Statistical properties of the state function *S*(*t*)

##### 2.1.1 Basic definitions

We consider a DNA-PAINT experiment with a single docking strand and introduce a state function *S*(*t*), which is 1 if the docking strand has bound an imager strand (‘on’-state) and 0 if it is vacant (‘off’-state).

The target switches from the state ‘0’ to the state ‘1’ with the association rate 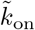, i.e. the 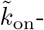 denotes the number of binding events per unit time. Note that the effective association rate 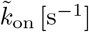 is defined as *k*_on_*c*_i_, with *k*_on_ [M^*−*1^s^*−*1^] the molar association rate and *c*_i_ [M] the imager concentration. The opposite process, ‘1’ to ‘0’, happens with the dissociation rate *k*_off_ . We assume that both rates are constant during an experiment. From the main text, 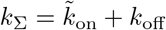 and 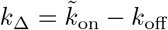 .

In an experiment, the state of the system is typically sampled at *n* equidistant time points separated by time-lapse Δ*t*, i.e., *t*_*j*_ = *j*Δ*t, j* = 1, 2, …, *n*, and *S*(*t*_*j*_) = *S*_*j*_. The discrete Fourier transform of the signal *S*_*j*_ is defined as

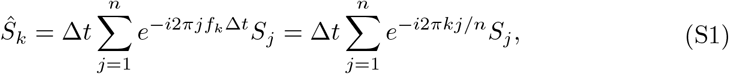

where *k* = 0, 1, …, *n* − 1, and we introduced the frequency

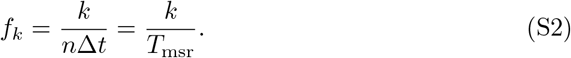

#### 2.1.2 Time evolution

We let *P*_0_(*t*) and *P*_1_(*t*) denote the probability that the system is in the state ‘0’ or ‘1’ at time *t*, respectively. The master equations that describe the time evolution of the system are

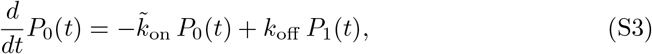

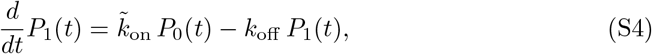

which are supplemented by the boundary conditions *P*_0_(*t*_0_) and *P*_1_(*t*_0_) for the state of the system at the initial time *t*_0_, and the normalization constraint *P*_0_(*t*) + *P*_1_(*t*) = 1 at all times.

These coupled, linear, first-order differential equations are solved using standard techniques, and the results are

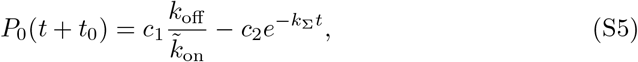

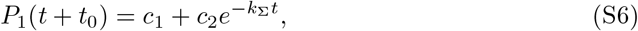

where the constants *c*_1_ and *c*_2_ are fixed by the boundary conditions *P*_0_(*t*_0_) and *P*_1_(*t*_0_).

For example, if the system is in state ‘0’ at time *t*_0_, then 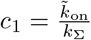 and 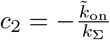 . So, the probability to still be in state ‘0’ at time *t* + *t* is 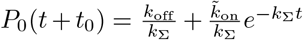, and the probability to be in state ‘1’ is 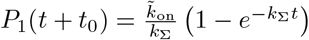.

##### 2.1.3 Transition probabilities

We now find the probabilities that the system is either in the same state or has transitioned to the other state in the time interval from 0 to *t*. For example, for the transition from the state ‘0’ to ‘1’, we denote this transition probability *P*_0*→*1_(*t*). The transition probabilities are,

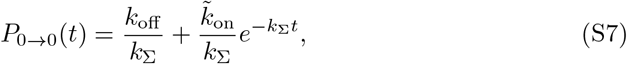

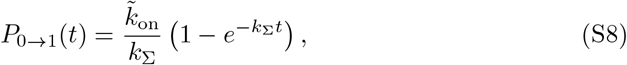

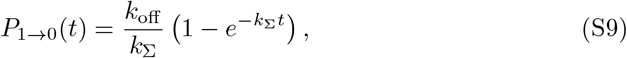

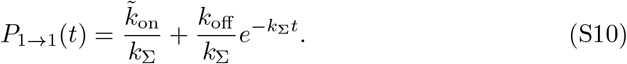

For *k*_Σ_*t* ≫ 1 holds that both *P*_0*→*0_(*t*) and *P*_1*→*0_(*t*) tend to

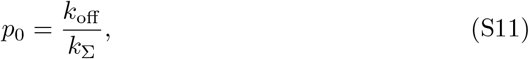

while *P*_0*→*1_(*t*) and *P*_1*→*1_(*t*) tend to

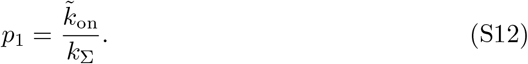

That is, the probabilities that the system is in state ‘0’ or ‘1’ at a random time point are *p*_0_ and *p*_1_, respectively. Notice the normalization condition, *p*_0_ + *p*_1_ = 1.

##### 2.1.4 Mean, variance, and third-order cumulant

The expected value of state function *S*(*t*) is

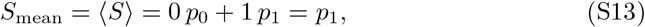

the variance is

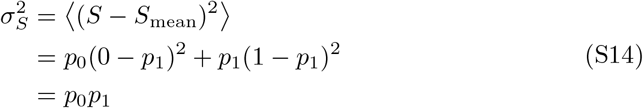

and the third-order cumulant is

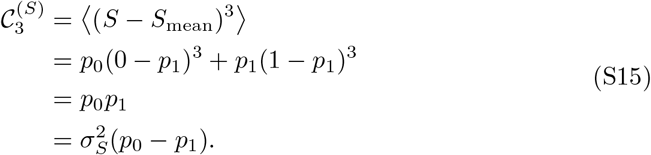

Inserting the expression for *p*_0_ and *p*_1_ from Eqs. (S11, S12) gives the results stated in the main text for the cumulants (see the section ‘Model parameters from the cumulants of a DNA-PAINT intensity trace’).

##### 2.1.5 Second-order correlation functions

The two-point correlation *ϕ*_2_(*t, t*^*′*^) is the expected value of the product between the intensity at two different time points, *t* and *t*^*′*^. With the transition probabilities defined above we get for *t*^*′*^ > *t* [2]

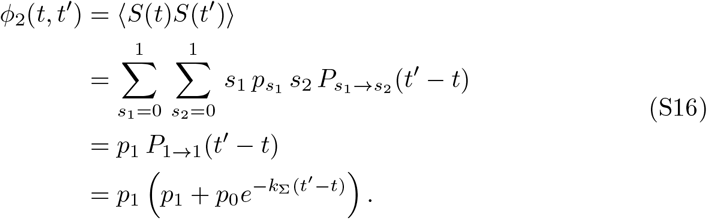

After rearranging and also considering *t*^*′*^ < *t*, the two-point correlation function becomes

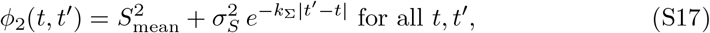

where | · | means absolute value.

The second-order cumulant function, also known as the autocovariance, is

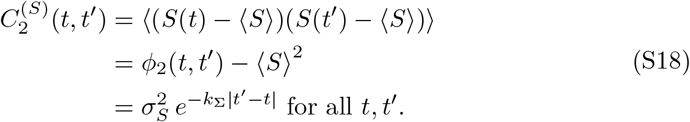

##### 2.1.6 Continuous and discrete power spectra

According to the Wiener-Khinchin theorem, the power spectrum is the Fourier transform of the autocovariance [2, 3]

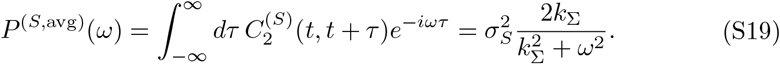

The discrete power spectrum (also named the periodogram) is defined as

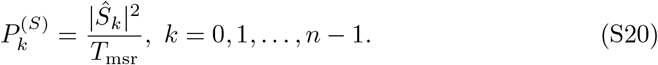

The average values of the power spectral values are (see derivation in Sec. 2.2)

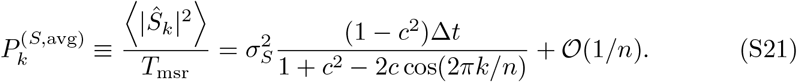

Equation (S21) includes the effect of aliasing due to the finite sampling rate [4].

##### 2.1.7 Third-order correlation functions

The three-point correlation function *ϕ*_3_(*t, t*^*′*^, *t*^*′′*^) is the expected value of the product between the intensity at three different time points, *t, t*^*′*^, and *t*^*′′*^.

With the transition probabilities defined above, we get for *t* < *t*^*′*^ < *t*^*′′*^,

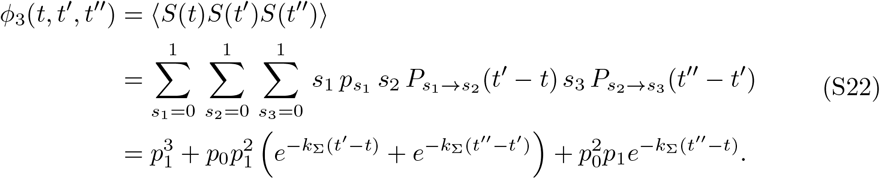

as only the term with (*s*_1_, *s*_2_, *s*_3_) = (1, 1, 1) is non-vanishing. Similar expressions are derived for the other permutations of the time points.

The third-order cumulant function of *S*(*t*) is

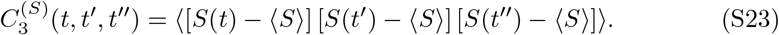

The time points can be ordered in six different ways. We here only show the three permutations needed for the derivation of the bispectra.

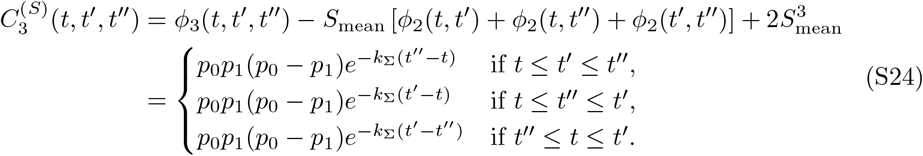

##### 2.1.8 Continuous and discrete bispectra

The continuous bispectrum is the two-dimensional Fourier transform of 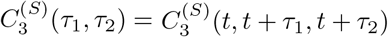

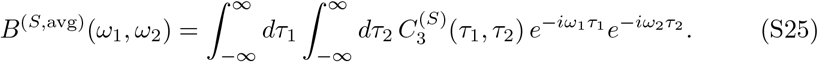

The derivation is shown in Sec. 2.3, and the results is (*ω*_3_ = *ω*_1_ + *ω*_2_)

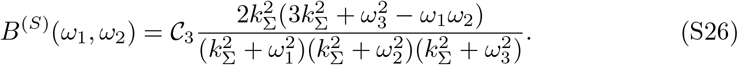

The bispectrum for discretely sampled intensities is derived in Sec. 2.4, and the result is (*k*_3_ = *k*_1_ + *k*_2_)

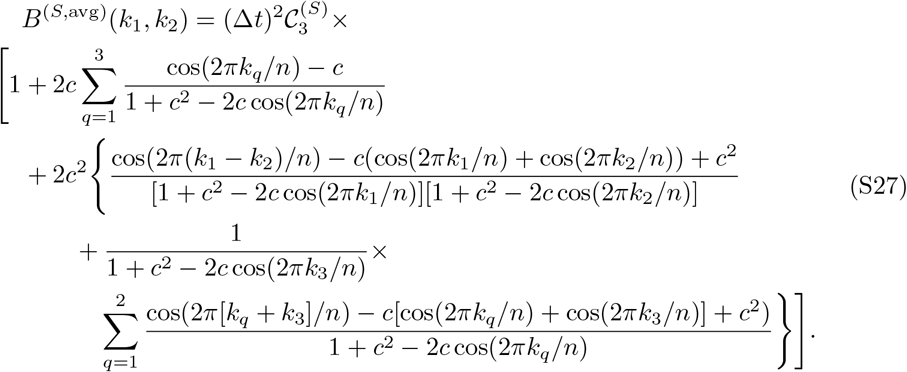

#### 2.2 Derivation of the discrete power spectrum

The expected value of the modulus-squared Fourier coefficients in Eq. (S1) gives

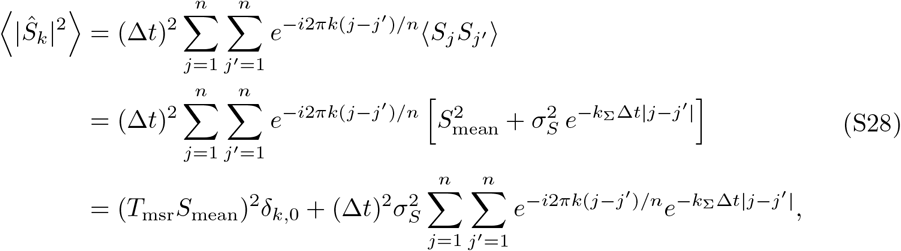

where we used that the first term in the bracket is independent of *j* and *j*^*′*^. Considering only finite frequencies, we get for *k* = 1, 2, …, *n* − 1 that

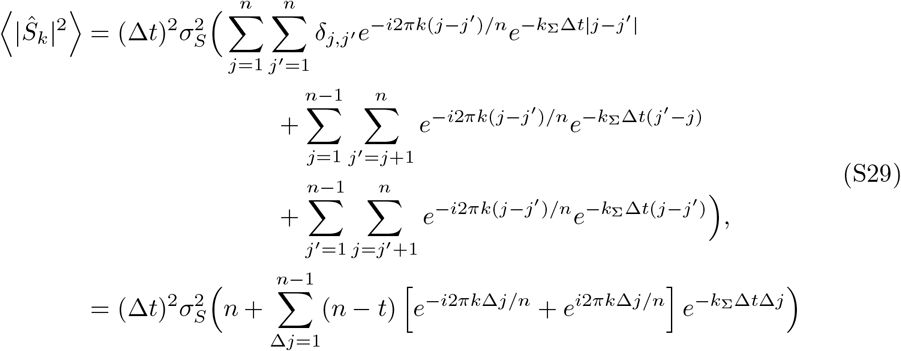

where we introduced Δ*j* = *j* − *j*^*′*^ in the last step.

The finite geometric series can be solved analytically, and to order *n* we find^1^

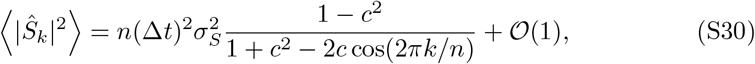

where 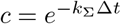 as defined in the main text.

Inserting this in the expression for the discrete power spectrum in Eq. (S21) gives to order 1*/n*

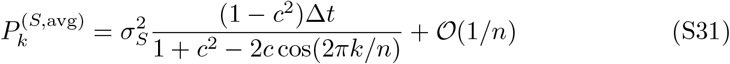

#### 2.3 Derivation of the continuous bispectrum

The continuous-frequency bispectrum is defined as the two-dimensional Fourier transformation of the third-order cumulant function [see Eqs. (S23,S24)]

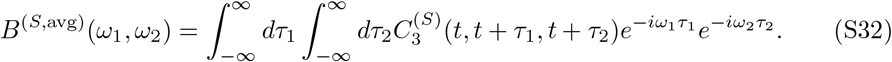

with *ω*_*q*_ = 2*πf*_*q*_ (*q* = 1, 2), where *f*_*q*_ is the frequency.

We split the integrals into the six segments, i) 0 < *τ*_1_ < *τ*_2_, ii) 0 < *τ*_2_ < *τ*_1_, iii) *τ*_2_ < 0 < *τ*_1_, iv) *τ*_1_ < 0 < *τ*_2_, v) *τ*_1_ < *τ*_2_ < 0, and vi) *τ*_2_ < *τ*_1_ < 0.

For i), we get

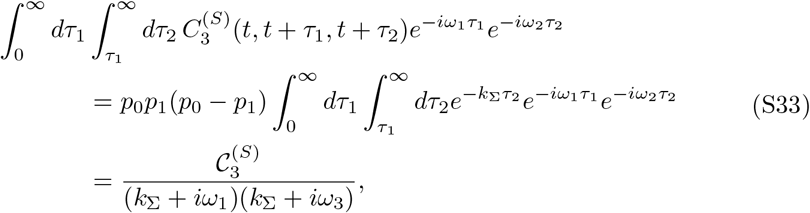

where we used the definition of 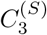 and Eq. (S24) in the second line, used 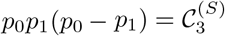, and introduced *ω*_3_ = *ω*_1_ + *ω*_2_.

For ii),

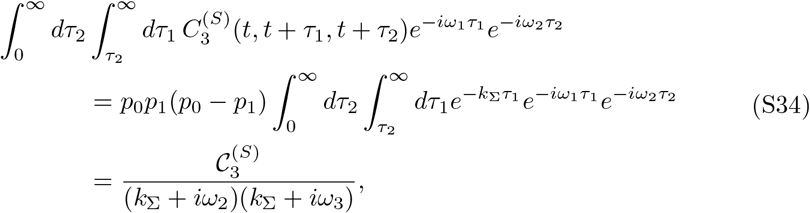

and finally for iii),

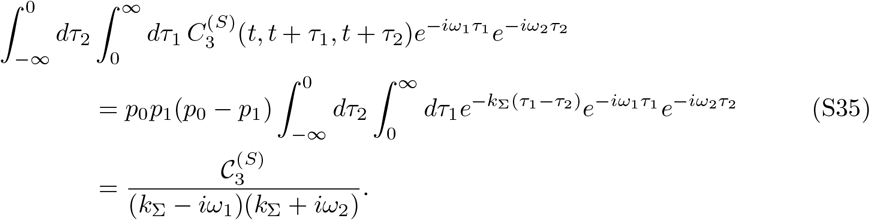

The other three segments give the complex conjugates of these expressions.

The sum of the integrals from all six segments gives the final expression for the continuous-time bispectrum,

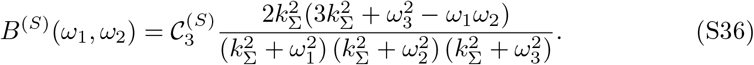

#### 2.4 Derivation of the discrete bispectrum

The discrete bispectrum is the (normalized) discrete Fourier transformation of the third-order cumulant,

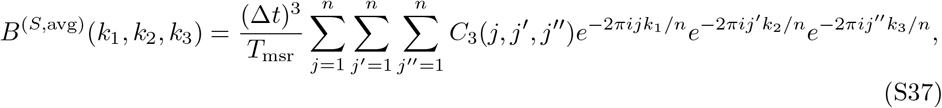

where *C*_3_(*j, j*^*′*^, *j*^*′′*^) ≡ *C*_3_(*t*_*j*_, *t*_*j*_*′, t*_*j*_*′′* ).

Due to time-translation invariance, the correlations can only depend on time differences. This implies that *k*_1_ + *k*_2_ + *k*_3_ = 0, such that

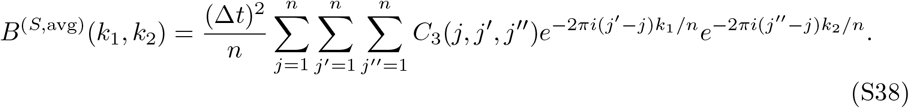

We now split the sum in Eq. (S38) into parts where all indices are different, two indices are identical, and all indices are identical. The resulting terms are different finite, geometric series, which we solve using *Mathematica* [Wolfram Research, Inc., Champaign, IL]

We start by considering the case where all indices are different. For *j* < *j*^*′*^ < *j*^*′′*^,

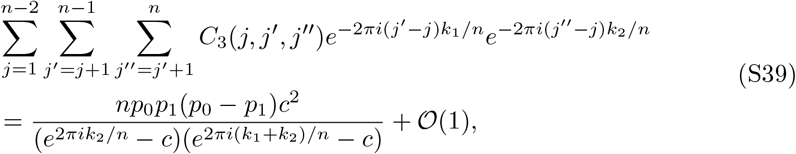

where 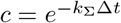 as defined in the main text.

If *j* < *j*^*′′*^ < *j*^*′*^, we get

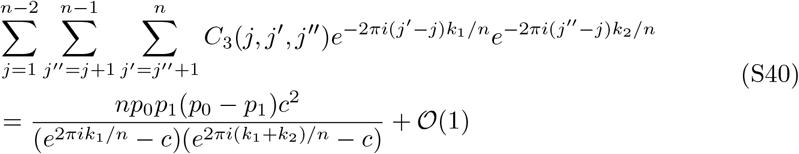

and finally for *j*^*′′*^ < *j* < *j*^*′*^,

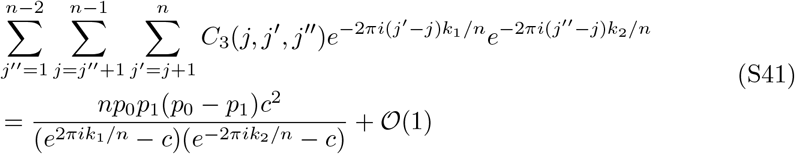

The other permutations of the indices give the complex conjugates of the terms stated above.

Now consider the case with two identical indices. For *j* = *j*^*′*^ < *j*^*′′*^,

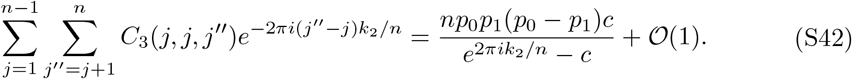

For *j* < *j*^*′*^ = *j*^*′′*^,

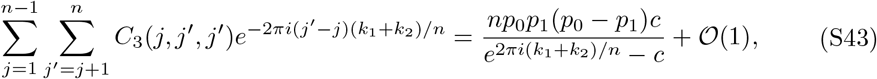

and, finally, for *j*^*′′*^ = *j* < *j*^*′*^,

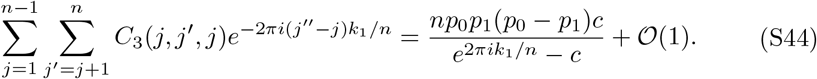

Again, the other permutations of the indices give the complex conjugates of the terms stated above.

When all of the indices are equal (*j* = *j*^*′*^ = *j*^*′′*^), the result is

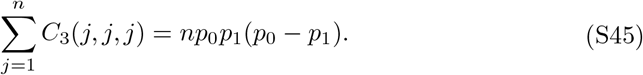

Adding up the terms and neglecting terms of order 𝒪(1*/n*) gives the normalized bispectrum,

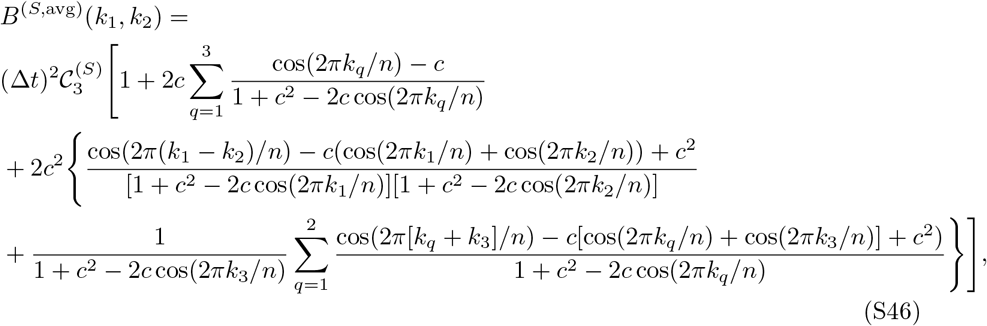

with *k*_3_ = *k*_1_ + *k*_2_.

#### 2.5 Simulating a trajectory of the state function *S*(*t*)

We here outline an efficient and numerically exact algorithm for simulating the state function of a molecular assembly with *N* independent targets, all characterized by the same constant kinetic parameters 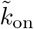 and *k*_off_ .

As the rates 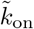 and *k*_off_ are zero-order reaction rates, i.e. they are constant, the waiting times between events are exponentially distributed (by definition of a Poisson process). So, if binding happens with a fixed rate 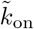, the time in the dark state is exponentially distributed with expected value 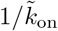,

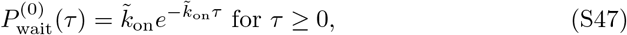

Similarly, the duration of time intervals in the bright state (imager bound) are exponentially distributed with expected value 1*/k*_off_,

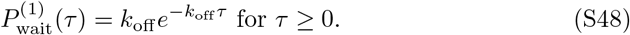

Hence we can simulate a realization of *S*(*t*) for a single target as follows: Choose an initial state according to the probabilities *p*_0_ and *p*_1_. As an example, assume that the system is in state ‘0’ at the initial time *t*_0_, *S*(*t*_0_) = 0. Then draw a random number *τ* ^(0)^ from the waiting time distribution in Eq. (S47), and update the time to 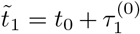 and the state to 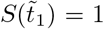. Then draw a new random number 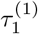 from the waiting time distribution in Eq. (S48), update the time to 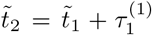, and the state to 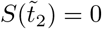. Continue until the length of the trajectory exceeds the desired measurement time *T*_msr_.

The algorithm determines the time points 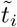 where the target changes state. The state of the system at the equidistant time points *t*_*j*_ = *j*Δ*t* sampled in an experiment is 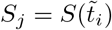, where 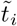 is the largest time-point for which it holds that 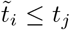. In other words, the value of the state function at discrete time point *t*_*j*_ takes the value of the last target state before that moment in time.

The outlined procedure is carried out independently for all *N* targets, and the total state of the molecular assembly 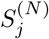 is is the sum of the states for the individual targets.

## Footnotes

1 We do not have to do this approximation, but do it to keep formulas as simple as possible.

